# Accurate forecasts of the effectiveness of interventions against Ebola may require models that account for variations in symptoms during infection

**DOI:** 10.1101/592030

**Authors:** W.S. Hart, L.F.R. Hochfilzer, N.J. Cunniffe, H. Lee, H. Nishiura, R.N. Thompson

## Abstract

Epidemiological models are routinely used to predict the effects of interventions aimed at reducing the impacts of Ebola epidemics. Most models of interventions targeting symptomatic hosts, such as isolation or treatment, assume that all symptomatic hosts are equally likely to be detected. In other words, following an incubation period, the level of symptoms displayed by an individual host is assumed to remain constant throughout an infection. In reality, however, symptoms vary between different stages of infection. During an Ebola infection, individuals progress from initial non-specific symptoms through to more severe phases of infection. Here we compare predictions of a model in which a constant symptoms level is assumed to those generated by a more epidemiologically realistic model that accounts for varying symptoms during infection. Both models can reproduce observed epidemic data, as we show by fitting the models to data from the ongoing epidemic in the Democratic Republic of Congo and the 2014-16 epidemic in Liberia. However, for both of these epidemics, when interventions are altered identically in the models with and without levels of symptoms that depend on the time since first infection, predictions from the models differ. Our work highlights the need to consider whether or not varying symptoms should be accounted for in models used by decision makers to assess the likely efficacy of Ebola interventions.

## 1. INTRODUCTION

Ebola epidemics have devastating consequences. The current epidemic in the Democratic Republic of Congo is the second largest in history, with 663 cases (614 confirmed and 49 probable) having been recorded as of 15^th^ January 2019 [1]. Mathematical models are increasingly used for exploring the effects of different possible control interventions during Ebola epidemics [2–4]. The values of model parameters are chosen so that the model output matches observed epidemic data (model fitting; Fig 1A), and then interventions are introduced in the fitted model to predict how the course of the epidemic is altered by different possible control strategies (intervention testing; Fig 1B).

**Figure 1.**
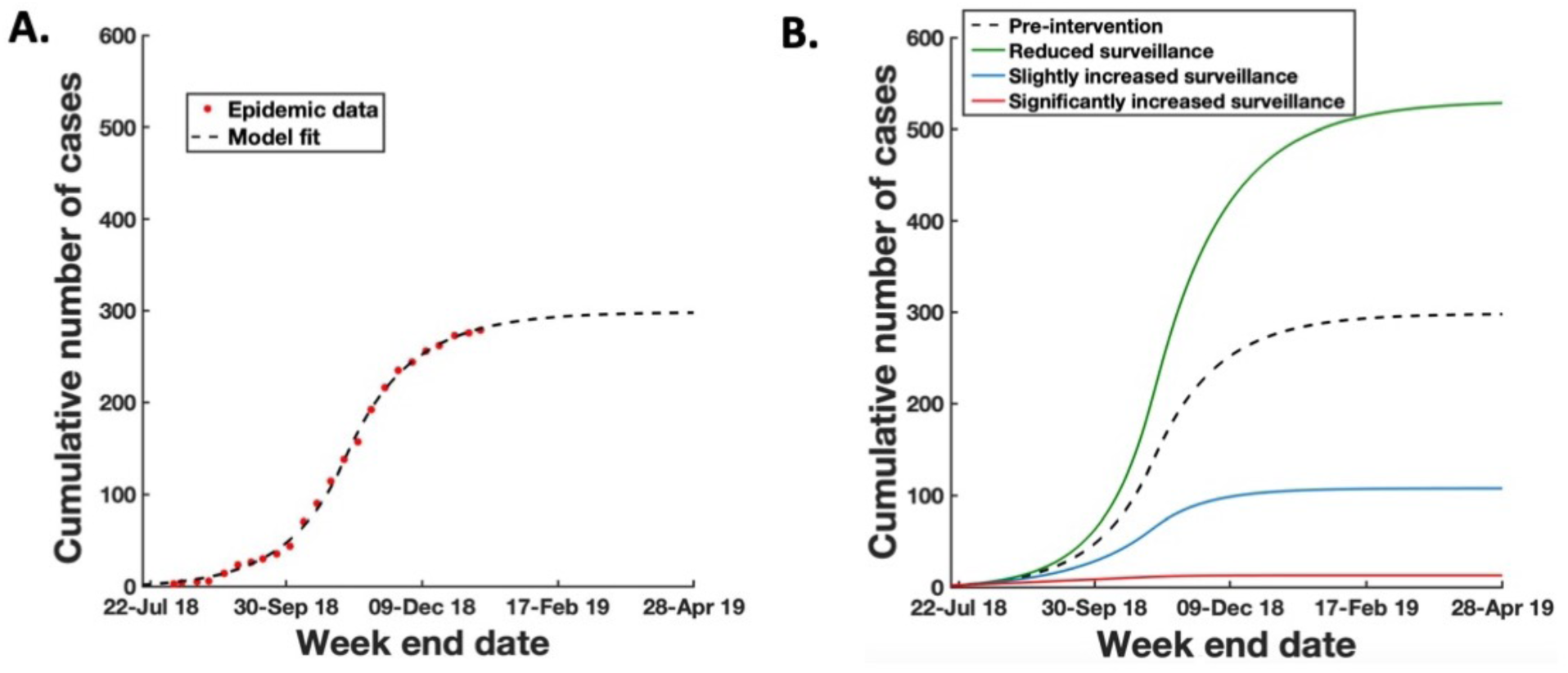
Schematic showing how a model can be used to predict the effect of changing control interventions on epidemic dynamics. Here, an intensification of surveillance is assumed to lead to improved detection and control of infectious hosts, thereby reducing the total number of cases. A. Model fitting. Model parameters are chosen so that the model output (black dotted) approximates epidemic data (red stars) B. Intervention testing. A range of alternative control interventions are introduced into the fitted model, and predicted dynamics under these new control interventions can be observed – predictions of the effects of reduced surveillance (green), slightly intensified surveillance (blue) and significantly intensified surveillance (red).

A commonly used model for characterising epidemics of diseases including Ebola is the Susceptible-Exposed-Infectious-Recovered (SEIR) model [5–7], and extensions to this basic model include explicit incorporation of transmission from Ebola deceased hosts [8–10] or accounting for mismatches between symptoms and infectiousness [11,12]. Possible interventions include isolation of symptomatic hosts, which can be included in the SEIR model by removing individuals from the infectious class. All individuals in the infectious class are usually assumed to be symptomatic, with the level of symptoms being assumed constant and therefore independent of the stage of infection. As an example, Chowell *et al.* [13] assume that symptomatic individuals are isolated at a constant rate, and Meakin *et al*. [4] assume that symptomatic individuals are hospitalised at a constant rate.

However, in reality, it is not the case that all symptomatic hosts are equally symptomatic. During an Ebola infection, an infected host progresses through different stages [14] – from initial non-specific symptoms (fever, headache and myalgia) to a gastrointestinal phase (diarrhoea, vomiting, abdominal symptoms and dehydration), and then either to a deterioration phase (collapse, neurological manifestations and bleeding) or recovery. Individuals with non-specific symptoms are less likely to be observed and treated/isolated than individuals who have progressed further through infection and have developed more specific and more serious symptoms.

Here, we investigate whether explicitly accounting for variations in symptom expression during the course of an Ebola infection leads to different epidemiological model dynamics compared to assuming a constant level of symptoms. To do this, we compare predictions derived from a model in which infectious hosts have a constant level of symptoms (the constant symptoms model – see Methods) with those from a model in which variable symptoms during infection are accounted for (the variable symptoms model). We parameterise our models using data from the ongoing Ebola epidemic in the Democratic Republic of Congo. We find that both models can be fitted closely to data from the epidemic. However, when control interventions in the models are altered, for example to explore the effects of intensifying surveillance and control, forecasts generated by the models are very different. We find the same qualitative result when we instead parameterise our models using data from the largest Ebola epidemic in history: the 2014-16 epidemic in west Africa.

These analyses demonstrate that models with or without variable symptoms can reproduce observed disease incidence time series, but that predictions from the models are different when interventions are altered, even when the change in interventions is identical in both models. Our results highlight the need to consider whether variations in symptom expression during infection should be included in models of Ebola epidemics. Without accounting for variable symptoms, predictions of the possible effects of interventions may be incorrect.

## 2. METHODS

### Datasets

To show that our results are not conditioned on particular properties of data from a single epidemic, we conducted two separate analyses in which we considered data from two different Ebola epidemics.

In the first analysis, we used data on the numbers of cases in approximately weekly time intervals from the ongoing Ebola epidemic in the Democratic Republic of Congo. It has recently been suggested by Dr Peter Salama, Deputy Director-General of Emergency Preparedness and Response at the World Health Organization, that this epidemic comprises several distinct outbreaks in different affected areas. Indeed, disease incidence time series display distinct phases (large numbers of cases at the end of July/beginning of August 2018, followed by low numbers of cases in September, and then larger numbers of cases again thereafter), probably due to spatial effects of spread of the virus which are not captured by standard non-spatial compartmental models [15]. For this reason, we focussed on data from the health zone of Beni, a city in the north-east of the Democratic Republic of Congo, and the neighbouring health zone of Kalunguta. This region has been severely impacted by the current epidemic. These data were obtained from World Health Organization disease outbreak news reports from 4^th^ August 2018 to 10^th^ January 2019 (Data S1, see also [16]).

In the second analysis, we considered data comprising of the numbers of cases in approximately weekly time intervals in Liberia during the 2014-16 Ebola epidemic, which were obtained from the World Health Organization (Data S2, see also [17]).

### Mathematical model

In the commonly used SEIR model, individuals are classified according to whether they are (*S*)usceptible to infection, (*E*)xposed, (*I*)nfected by the pathogen or (*R*)emoved and no longer infectious. We extended this model to account explicitly for case finding followed by isolation of infectious individuals. We also assumed that there were three distinct phases of infection, corresponding to different stages of an Ebola infection. This delivered the additional benefit that the infectious period (in the absence of control) was gamma distributed, rather than exponentially distributed – and gamma distributions have been found to characterise epidemiological periods accurately in a range of systems [18,19]. This gave rise to the SEI_1_I_2_I_3_RC model,

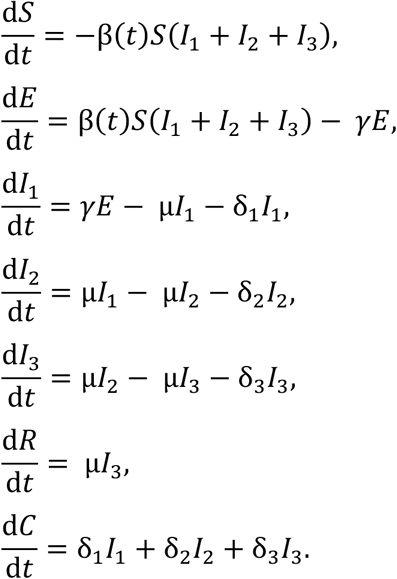

In this model, the *C* compartment represents the number of individuals that have ever been controlled (detected and isolated) until the current time. Since we wish to isolate the impacts on prediction of variable symptoms alone, in the baseline version of the model we assume that all infectious hosts are equally infectious – although we consider the effect of relaxing this assumption later.

In our analyses, we made the assumption widely used in Ebola models that the infection rate parameter in the model is temporally-varying [4,5], to reflect changes in transmissibility during the epidemic. This could, for example, indicate changes in behavioural responses or alterations to interventions (aside from detection and isolation, since we model that explicitly). In particular, we assumed that

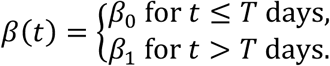

We then considered two alternative versions of the model. In the first (the constant symptoms model – Fig 2A), we assumed that all infectious individuals are successfully detected and isolated at the same average rate per day, so that δ_1_ = δ_2_ = δ_3_ = δ, say. This assumption is common to any epidemiological model that includes interventions aimed at symptomatic hosts, unless differences in symptom expression are accounted for explicitly. The constant symptoms model is therefore similar to most epidemiological models that have been used to represent Ebola epidemics previously (e.g. [4,13]).

**Figure 2.**
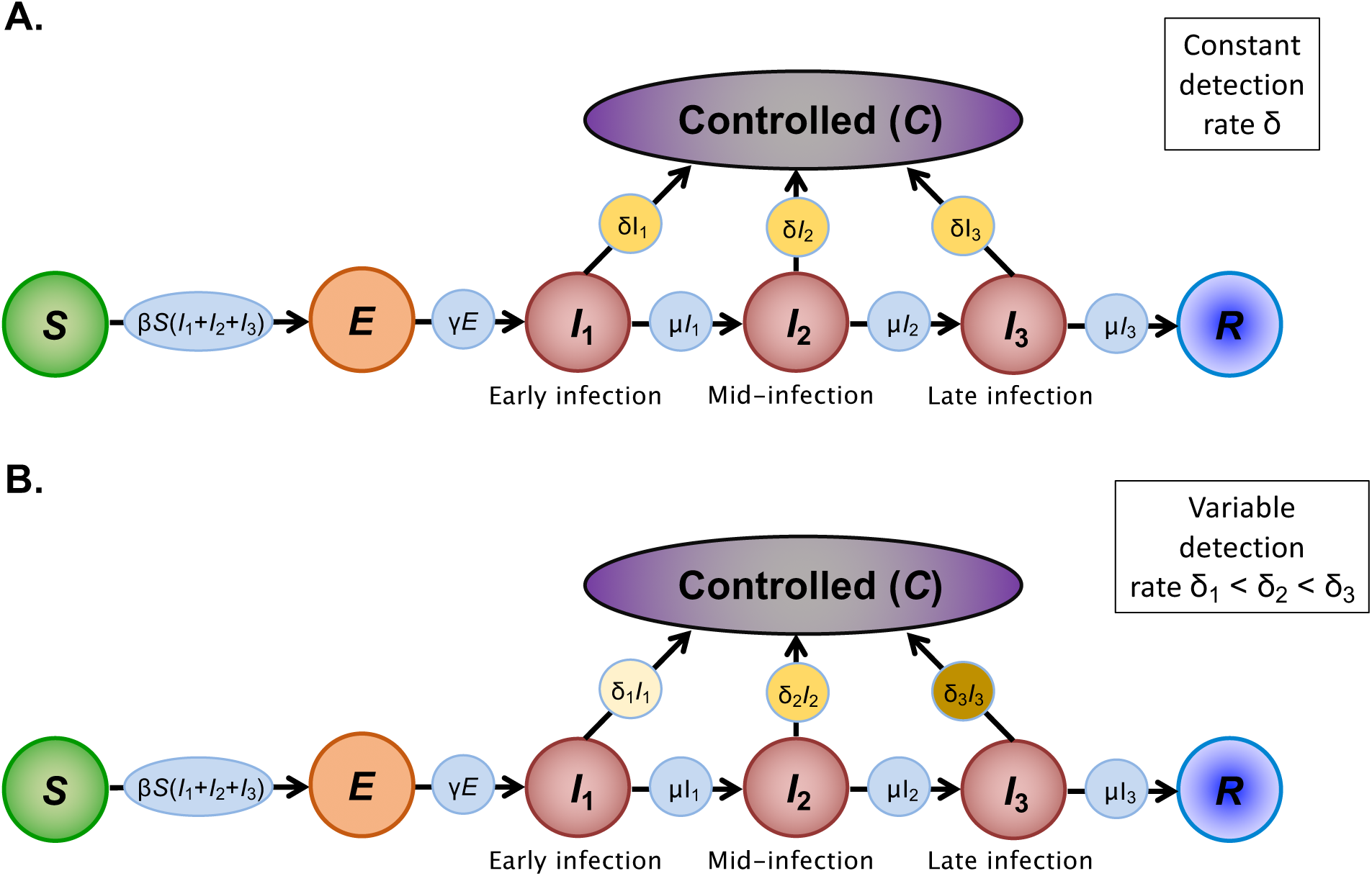
Schematic of the different models that we considered. A. Constant symptoms model, in which individuals in each stage of infection are equally likely to be detected and isolated (so that the detection rate, δ, is equal for all three infectious classes); B. Variable symptoms model, in which symptoms are assumed to intensify during an Ebola infection (so that the detection rate is smaller for individuals in earlier infection compared to later infection, i.e. δ_1_ < δ_2_ < δ_3_). We also show how additional epidemiological complexity can be included in these models (see Supplementary Material).

We also considered the more realistic case in which symptoms become more severe as infection progresses, so that δ_1_ < δ_2_ < δ_3_. We refer to the resulting model as the variable symptoms model (Fig 2B). This model reflects the fact that, in reality, individuals with initial mild symptoms are less likely to be detected and isolated to prevent further transmission than individuals with more developed symptoms who are in the gastrointestinal or deterioration phases.

### Model fitting and parameters

We considered the numerical solutions of the models described above in a host population of size of *S* + *E* + *I*_1_ + *I*_2_+ *I*_3_+ *R* + *C = N* individuals and a basic reproduction number at the beginning of the epidemic and in the absence of surveillance given by 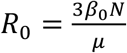.

The default parameter values used in our analyses are given in Table 1 (for the Democratic Republic of Congo in 2018-19) and Table 2 (for Liberia in 2014-16).

However, as described in the Results, we also checked the robustness of our results to these particular parameter values. The values of the infection rates *β*_0_ and *β*_1_, as well as the date on which the infection rate changes, *T*, were obtained by fitting the outputs of the models to the epidemic data. The start date of the epidemic, *T*_0_, was also estimated during the fitting procedure. Model fitting was performed using least squares estimation – i.e. choosing parameter values to minimise the sum of squares distance between the cumulative numbers of detected or removed hosts in the model (*C* + *R*) and the cumulative numbers of cases in the data. Numerical solutions were generated starting with a single host in the *E* compartment at the start time of the epidemic, *T*_0_, with all other individuals susceptible.

**Table 1.** Default parameter values used in our analysis of data from the ongoing Democratic Republic of Congo epidemic.

| Parameter | Definition | Default Value | Justification |
| --- | --- | --- | --- |
| $N$ | Population size | 230,000 | [42] |
| $\beta_0$ | Infection rate early in epidemic | $8.43 \times 10^{-7} \text{ day}^{-1}$<br>(constant symptoms model)<br>or $8.1 \times 10^{-7} \text{ day}^{-1}$<br>(variable symptoms model) | Fitted to data |
| $t_{\text{ay}}\beta_1$ | Infection rate late in epidemic | $3.68 \times 10^{-7} \text{ day}^{-1}$<br>(constant symptoms model)<br>or $3.59 \times 10^{-7} \text{ day}^{-1}$<br>(variable symptoms model) | Fitted to data |
| $T$ | Infection rate switch time | 25<sup>th</sup> October 2018<br>(constant symptoms model)<br>or 24<sup>th</sup> October 2018<br>(variable symptoms model) | Fitted to data |
| $1/\gamma$ | Latent/incubation period | 7 days | [4] |
| $1/\mu$ | Infectious period | 9.8 days | [4] |
| $p_1$ | Detection probability in early infection | 0.6 (constant symptoms model)<br>or 0.1 (variable symptoms model) | Assumption (for analyses with different values, see Supplementary Material) |
| $p_2$ | Detection probability in mid infection | 0.6 (constant symptoms model)<br>or 0.8 (variable symptoms model) | Assumption (for analyses with different values, see Supplementary Material) |
| $p_3$ | Detection probability in late infection | 0.6 (constant symptoms model)<br>or 0.9 (variable symptoms model) | Assumption (for analyses with different values, see Supplementary Material) |
| $\Delta$ | Sampling frequency | 21 days (weak surveillance) or 14 days (intensified surveillance) | Assumption (for analyses with different values, see Supplementary Material) |
| $\delta_1$ | Detection/isolation rate in early infection | $4.08 \times 10^{-2} \text{ day}^{-1}$ (constant symptoms model)<br>or $5.01 \times 10^{-3} \text{ day}^{-1}$ (variable symptoms model) | Calculated using values of $p_1$ and $\Delta$ (see Methods) |
| $\delta_2$ | Detection/isolation rate in mid infection | $4.08 \times 10^{-2} \text{ day}^{-1}$<br>(constant symptoms model)<br>or $6.35 \times 10^{-2} \text{ day}^{-1}$<br>(variable symptoms model) | Calculated using values of $p_2$ and $\Delta$ (see Methods) |
| $\delta_3$ | Detection/isolation rate in late infection | $4.08 \times 10^{-2} \text{ day}^{-1}$<br>(constant symptoms model)<br>or $7.79 \times 10^{-2} \text{ day}^{-1}$<br>(variable symptoms model) | Calculated using values of $p_3$ and $\Delta$ (see Methods) |
| $T_0$ | Start date of epidemic | 25<sup>th</sup> June 2018<br>(constant symptoms model and variable symptoms model) | Fitted to data |

**Table 2.**
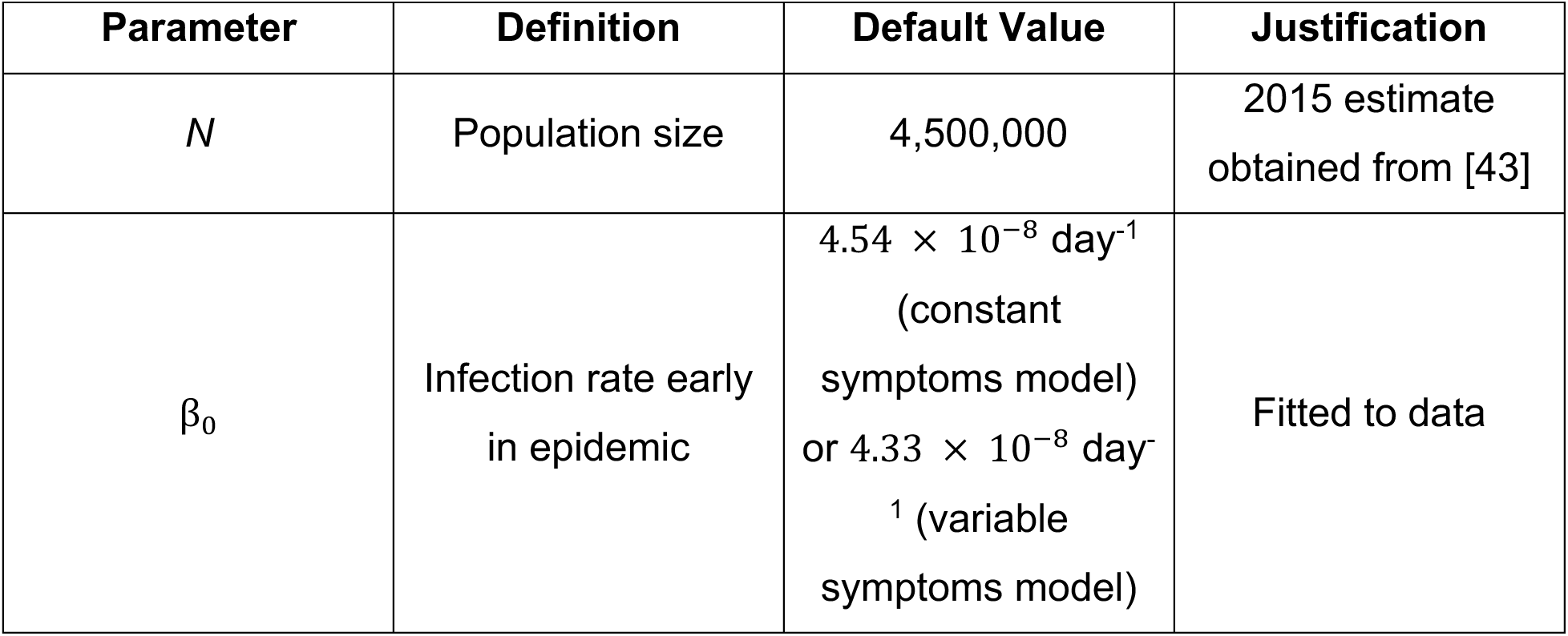
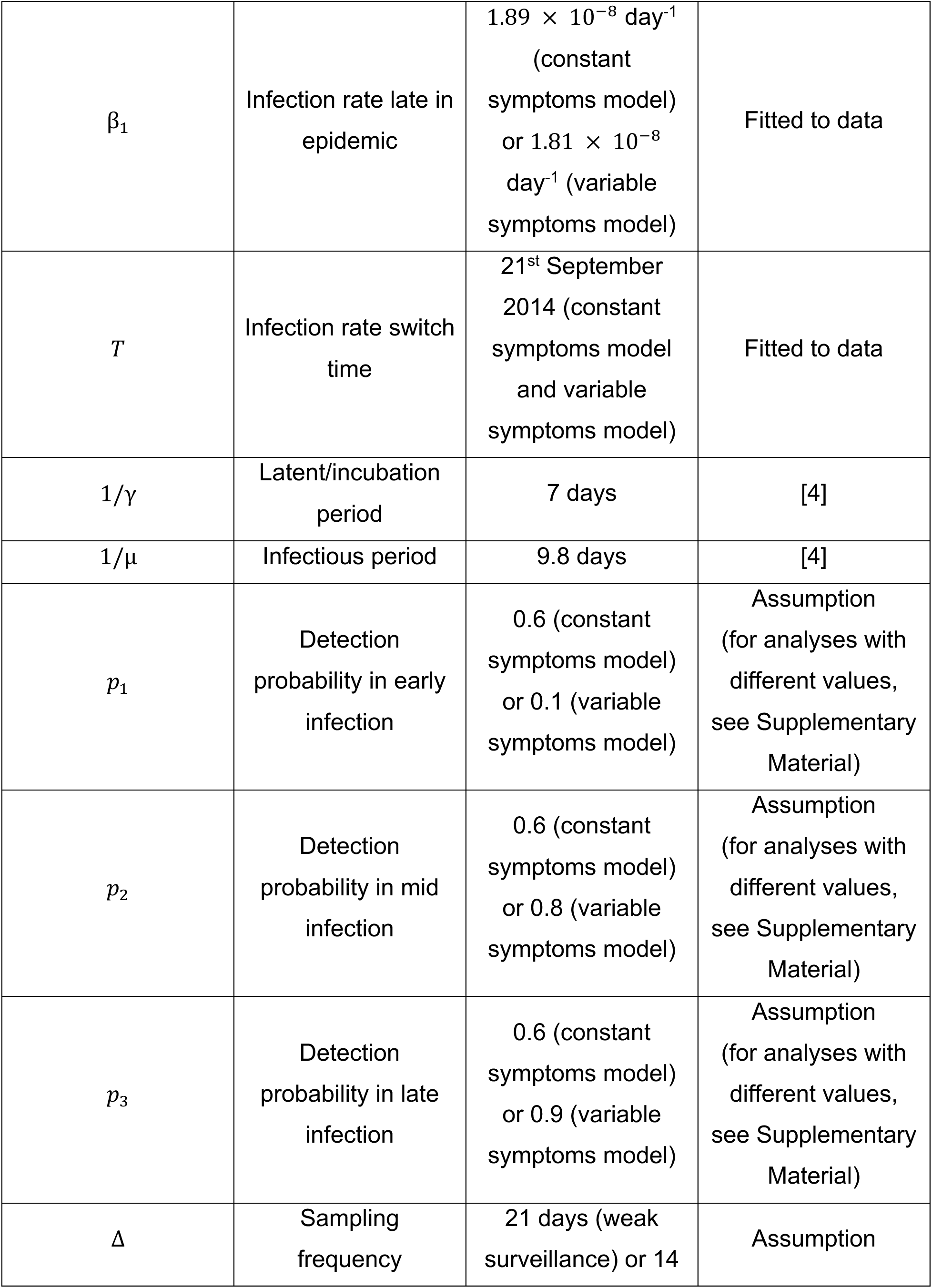
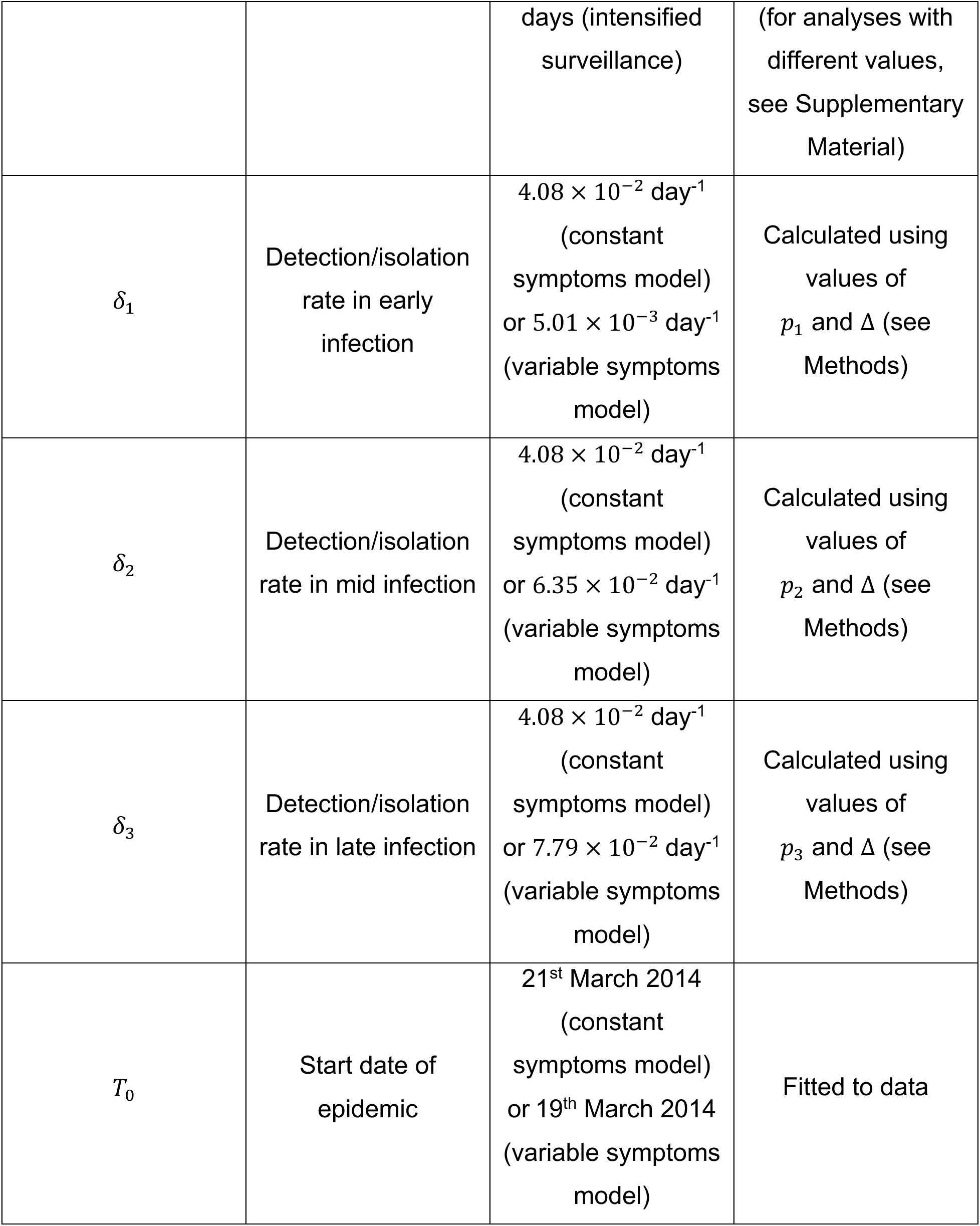
Default parameter values used in our analysis of data from the 2014-16 epidemic in Liberia.

The values of the parameters characterising the rate of Ebola detection and isolation, i.e. δ_1_, δ_2_ and δ_3_, depend on the level of surveillance, which includes various passive and active case finding strategies. We did not model explicitly the wide range of different surveillance activities that take place during an Ebola response (see Discussion). However, to provide a concrete setting in which to illustrate the principle that forecasts are different under the constant symptoms and variable symptoms models, we instead considered a simplified scenario in which each host is checked for infection on average every ∆ days. Each time monitoring occurs, there is a detection probability of *P*_*i*_ for individuals in class *I*_*i*_ (for *i*; = 1,2, or 3). As a result,

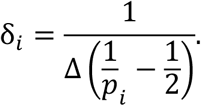

This expression is derived in the Supplementary Material (for a similar approach, see also [20]).

In our main analyses, we considered two different surveillance regimes. When the models were fitted, under weak surveillance, we assumed that the default surveillance period was ∆ = 21 days. When the fitted models were then used to predict the impacts of intensified surveillance, the surveillance period was changed to ∆ = 14 days. We assumed that the detection probability in the constant symptoms model was *P*_1_ = *P*_2_ = *P*_3_ = 0.6. When we accounted for the possibility that symptoms change as hosts progress through infection, we instead used default values of *P*_1_ = 0.1, *P*_2_ = 0.8 and *P*_3_ = 0.9 so that the mean value of *P*_1_, *P*_2_ and *P*_3_ was equal to the value of these parameters in the constant symptoms model. In other words, conditional on not being detected previously, a host chosen at a random time in the infectious period was equally likely to be detected in both models.

## 3. RESULTS

As described in the Introduction, fitted models are often used to test potential control interventions (Fig 1). We therefore considered fitting models to two different datasets – one from the current Ebola epidemic in the Democratic Republic of Congo, and another from the historical Ebola epidemic in west Africa in 2014-16.

First, we considered data from the ongoing Ebola epidemic in the Democratic Republic of Congo (Fig 3A). We fitted the constant symptoms model and variable symptoms model to these data in turn, and found that both of these models could replicate the observed dynamics of the epidemic (Figs 3B). We then used these fitted models to predict how the epidemic dynamics would have been altered under a different control intervention. In particular, we increased the rate of detection in the fitted models, to represent predictions under an intensification of surveillance and control efforts (see Methods).

**Figure 3.**
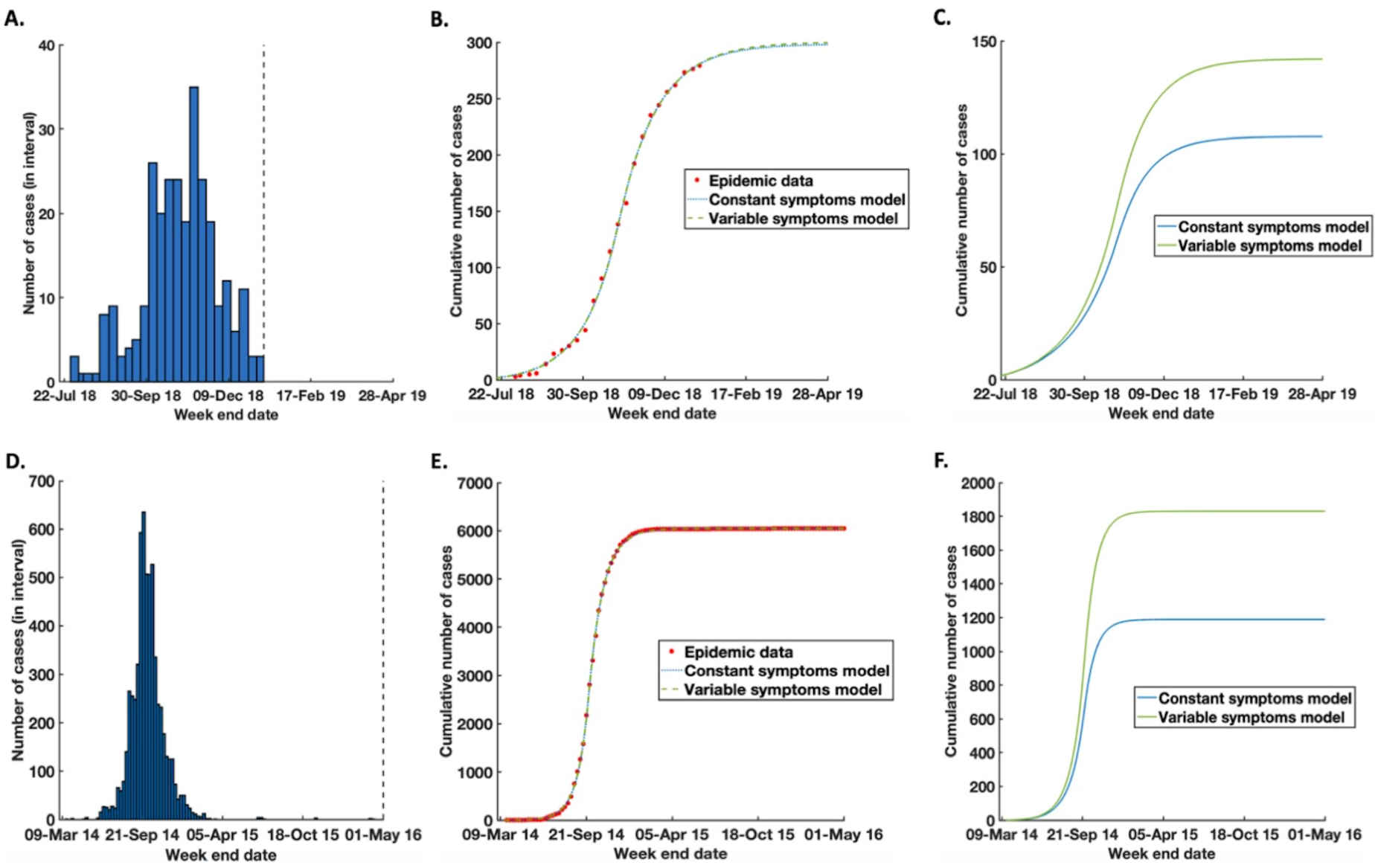
Using the constant symptoms model and variable symptoms model to predict alternative interventions. A. Number of cases each week in Beni and Kalunguta health zones in the Democratic Republic of Congo. The dotted black vertical line represents the time before which data were available. B. Model fits to the data (red stars), using the constant symptoms model (blue dotted) and variable symptoms model (green dash) C. Predictions of the effect of intensified surveillance, using the constant symptoms model (blue) and variable symptoms model (green). D-F. Equivalent figures to A-C, using the data from the 2014-16 Ebola epidemic in Liberia. For model parameters, see Methods and Tables 1 and 2.

When surveillance was intensified, the prediction of the constant symptoms model differed substantially from that of the variable symptoms model (Fig 3C). In particular, for the parameter values displayed here, the constant symptoms model predicted 24% fewer cases than the more epidemiologically realistic variable symptoms model (108 cases in the constant symptoms model as opposed to 142 cases in the variable symptoms model). Consequently, even though the observed dynamics of the models appear identical when fitted to data, they produce different predictions when control interventions are changed.

We then repeated our analysis, instead using data from the 2014-16 Ebola epidemic in west Africa (Figs 3D-F). In this case, the constant symptoms model predicted 35% fewer cases than the variable symptoms model (Fig 3F). Since the total number of cases in this epidemic was so large, this corresponded to 641 cases difference between the forecasts of the two models.

We also considered the robustness of our results to the values of the model parameters, including the level of surveillance assumed when fitting to data and the extent to which surveillance was intensified in the models (Figs S1-S7). In addition, we considered different values of the detection probability whenever surveillance occurs (Figs S8 and S9). In each case, we found qualitatively identical results – both the constant symptoms and variable symptoms models could reproduce epidemic time series data, but when interventions were changed in the models the predicted epidemic dynamics then differed. As well as considering an intensification of surveillance, we also examined cases in which surveillance was relaxed (e.g. Figs S4B and S4C). Whenever surveillance was intensified, the constant symptoms model underestimated the total number of cases compared to the more realistic variable symptoms model. However, when surveillance was instead reduced, the constant symptoms model led to overestimation of the total number of cases.

We also considered the effect of enhancing surveillance at different times during the epidemic. In particular, we considered increasing the surveillance level once the epidemics had already been in progress until a certain date, for different possible dates of surveillance intensification. The earlier that surveillance was intensified, the larger the error when using the constant symptoms model rather than the more biologically realistic variable symptoms model, since early surveillance intensification then allows more time for model predictions to differ (Figs 4 and S10).

**Figure 4.**
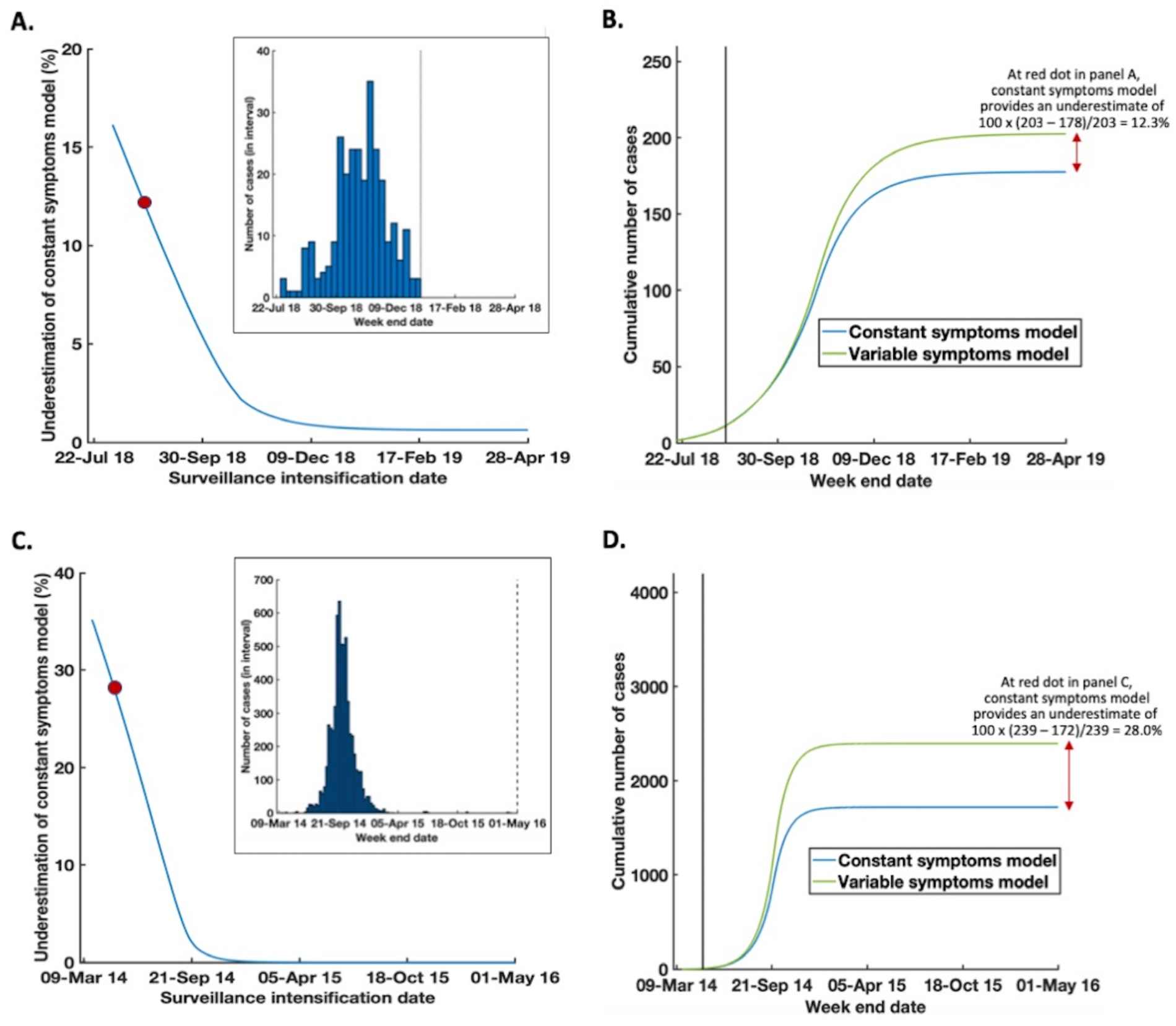
Using the constant symptoms model and variable symptoms model to predict alternative interventions, for different times of surveillance intensification. A. Reduction in the total number of cases predicted by the constant symptoms model compared to the variable symptoms model, expressed as a percentage of the number of cases predicted by the variable symptoms model, for different times of surveillance intensification. The models were fitted to data from the ongoing epidemic in the Democratic Republic of Congo. B. Illustration of how the values in panel A were calculated. In the graph shown, intensified surveillance began on 23^rd^ August 2018 in the models fitted to the ongoing epidemic in the Democratic Republic of Congo. This corresponds to the time denoted by the red dot in panel A. C-D. Equivalent panels to A-B, for the 2014-16 Ebola epidemic in Liberia. In panel D, intensified surveillance began on 2^nd^ May 2014. In panels A and C, we only consider intensifying surveillance at or after the time that the first cases were observed (i.e. 3^rd^ August 2018 for the ongoing epidemic in the Democratic Republic of Congo, and 23^rd^ March 2014 for the 2014-16 Liberia epidemic). The original epidemic datasets are shown as insets to panels A and C. For model parameters, see Supplementary Material and Tables 1 and 2.

Until this point, so that we could isolate the effect of variable symptoms alone on the predicted outcomes of interventions, we assumed that at any time during an epidemic all infected and uncontrolled hosts generated new infections at a constant rate. However, we also conducted an analysis in which the infection rate also varied throughout the course of an Ebola infection, by considering cases in which infectiousness was either correlated with or correlated against the level of symptom expression (Fig 5 and Supplementary Material). In cases in which higher levels of symptoms were associated with reduced infectiousness – for example due to a lower level of mixing in the population compared to hosts with less serious symptoms – our result that predictions are different between the constant symptoms and variable symptoms models was enhanced (Figs 5B and 5D).

**Figure 5.**
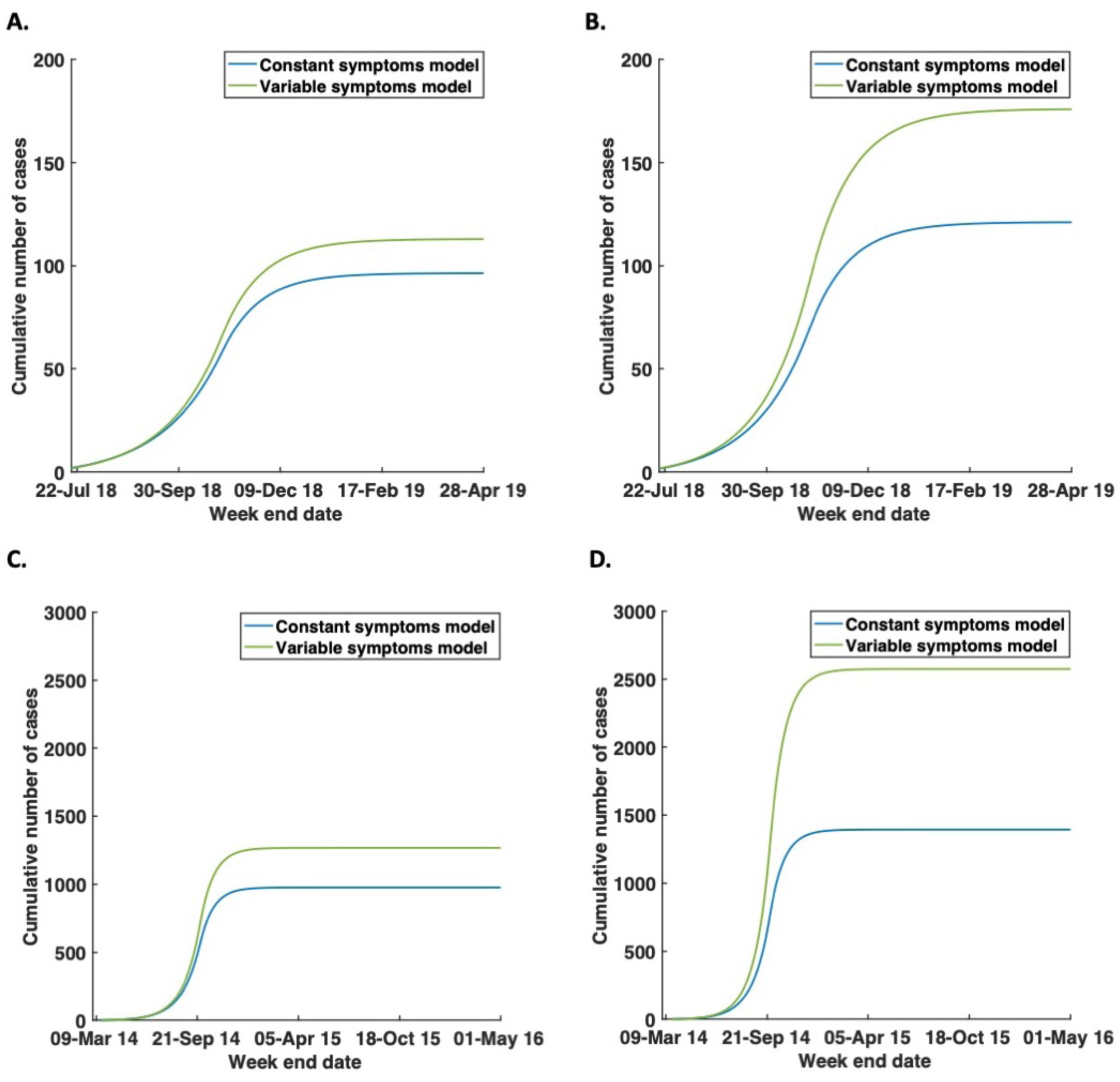
Using the constant symptoms model and variable symptoms model to predict alternative interventions, if infectiousness is also assumed to depend on the stage of infection. A. Predictions of the effect of intensified surveillance using the constant symptoms model (blue) and variable symptoms model (green), fitted to data from the ongoing epidemic in the Democratic Republic of Congo and assuming that infectiousness increases during an Ebola infection as described in the Supplementary Material. B. Predictions of the effect of intensified surveillance using the constant symptoms model (blue) and variable symptoms model (green), fitted to data from the ongoing epidemic in the Democratic Republic of Congo and assuming that infectiousness decreases during an Ebola infection as described in the Supplementary Material. C-D. Equivalent figures to A-B but fitted to data from the 2014-16 Ebola epidemic in Liberia. The values of parameters other than the fitted parameters (i.e. the infection rates, the start times of the epidemics and the times at which the infection rates change), are identical to those described in Tables 1 and 2. Surveillance intensification is assumed to occur at the beginning of the epidemic. Model fits are not shown, since they appear identical by eye to those in Figs 3B and 3E.

## 4. DISCUSSION

Epidemiological models that have been fitted to data are often used to predict the dynamics of an epidemic under different control interventions (e.g. [21–25]). However, most models considering interventions targeted at symptomatic hosts assume that those hosts display a constant level of symptoms that does not change during the course of infection. For a number of infectious diseases, however – including Ebola virus disease – there are different stages of an infection, and in each of these stages the level of symptoms is likely to be different. Individuals in early infection tend to have milder symptoms than those in later infection. As a result, symptomatic hosts in early infection are less likely to have appeared in the surveillance data that are routinely collected during an epidemic than those in later infection, who not only have had a longer period during which to be detected but are also likely to have developed more severe and recognisable symptoms.

Here, we have considered Ebola virus disease as a case study, and used two models to predict the possible effects on the dynamics of two epidemics under different surveillance levels. We assumed that increased surveillance leads to improved detection and control of symptomatic hosts. We compared the output of a model in which, if an individual is surveyed, the probability of detection is constant at each time during the infectious period (the constant symptoms model) to the equivalent predictions from a model in which the probability of detection increases throughout infection (the variable symptoms model). We found that both these models can be fitted closely to data from Ebola epidemics (Figs 3B and 3E). However, when the level of surveillance in the models is increased, we found that the more epidemiologically realistic variable symptoms model predicted a smaller number of cases (e.g. Figs 3C and 3F). Thus, it might be important to use the more realistic model to assess the quantitative effects of interventions aimed at reducing the impacts of Ebola epidemics.

Variations in symptoms between different stages of infection, as well as the signature of such variable symptoms in those types of data that are collected during an epidemic, have to date received little attention. Until now, the impact of variable symptoms on predictions of models used for testing Ebola interventions has never been rigorously assessed. However, our approach of splitting the infectious and symptomatic period into different compartments was inspired by the so-called “method of stages” [18,26], a technique most often used to model gamma distributed epidemiological periods [11,27,28]. Within that framework, varying infectiousness – rather than symptoms – over the course of infection has been considered previously. For example, Cunniffe *et al.* [19] consider a model of plant disease epidemics in which the rate of sporulation (production of viable spores by each infected host) is a function of the time since infection, and implement this in an SEIR model by splitting the *E* and *I* classes into compartments and assigning different infection rates to hosts in the different *I* classes. A similar modelling framework could be adopted in our work, using a large number of compartments so that the level of symptoms is represented by a continuous curve (rather than being at constant levels within the different stages of infection). However, we do not pursue this here, since discrete changes in symptom expression in each symptomatic host are sufficient to make our underlying point that accurate forecasts of the effects of Ebola interventions may require models that account for variations in symptoms.

To conduct our analyses, we sought to develop a simple model in which the level of symptoms increases during infection, and to compare the results from this model to those from the analogous model in which there is a constant level of symptoms during infection. Practical use of either model during an Ebola epidemic would require adjustment for the particular epidemic under consideration. For example, transmission in different settings could be included in a single model, such as spread in hospitals, community care centres, at funerals or in the wider community [3,4]. We modelled detection and isolation of symptomatic hosts here, but other interventions such as vaccination could be modelled explicitly [29]. If an Ebola vaccine is not perfectly effective, as has been suggested for the vaccine used in the ongoing Ebola epidemic [30], the possibility that vaccination might mask symptoms while not completely stopping infectiousness could be included in our approach. A model that includes spatial spread of the pathogen or transmission through social contact networks might be required to replicate observed data [31,32], or different geographical areas could be considered separately [2,4]. To demonstrate the principle that variable symptoms can affect predictions of the effects of interventions, we assumed that all infected individuals pass through three stages of an Ebola infection (from non-specific symptoms, to a gastrointestinal phase and then to a deterioration phase), whereas in reality some hosts might recover rather than passing to the deterioration phase [14]. At the cost of an additional parameter to be estimated, it would be straightforward to include this in a compartmental epidemiological model (see preliminary analysis in Supplementary Material and Fig S11, in which some hosts recover rather than passing to the final stage of infection). We also modelled surveillance in a simple fashion, by assuming that hosts are surveyed on average at periodic intervals and that there is a particular probability of detection whenever a host is surveyed. For forecasting, it would be necessary to consider the wide range of different surveillance approaches used in practice including contact tracing from known cases [33] and rural village visitations to detect cases in locations where access to healthcare is limited [34], as well as disruptions to surveillance caused by factors including armed conflict [35].

We parameterised our models using the simplest possible approach – namely fitting the numbers of detected or removed individuals in the relevant classes of the models to data on the cumulative numbers of symptomatic cases using least squares estimation. We did not quantify the uncertainty in estimates of the values of model parameters, since the precise method of parameter inference was not central to our message. Instead, we sought to use the simplest possible fitting method. While this approach is used frequently during epidemics due to its ability to produce quick forecasts [5,36,37], to properly quantify the uncertainty in forward projections it would be necessary to use non-cumulative incidence data and fit stochastic transmission models [38].

One advantage of the models that we used is that the surveillance level is assumed to impact on the epidemiological dynamics themselves, rather than simply the observed dynamics. This is not always the case in epidemiological models: a common method for accounting for under-reporting is simply to scale the incidence data up [39], thereby assuming a fixed percentage of infectious cases are detected with no impact on the numbers of cases generated by those individuals. Another approach is to assume that some individuals in the infectious class are unobserved [40]. In reality, detected hosts have a lower probability of transmitting the pathogen than undetected hosts due to the higher chance that those individuals are subject to interventions, and our models reflect this.

Here, we considered a control strategy of detection and isolation under different surveillance levels. The effects of including variable symptoms in models of other intervention strategies should be tested, to see whether it is always necessary to account for changing levels of symptoms throughout infection. We note that including additional epidemiological detail in forecasting models does not always improve predictions [12]. Simple models are easier to parameterise and interpret than more complex models, and so modellers should consider carefully, in each study, whether or not including variable symptoms will change model predictions. We also note that, in theory, it might be possible to deploy commonly used epidemiological models with altered parameter values as a proxy for explicit consideration of variable symptoms [41]. For example, if the chance of detection in early infection is low, then early non-specific symptoms could be considered as part of the incubation period. In that case, care should be taken when “lifting” the values of model parameters directly from the clinical literature, to ensure that the definitions of parameters in the model match those in the original studies.

In summary, including different levels of symptoms at different stages of infection in epidemiological models can alter predictions of the effects of intervention strategies compared to assuming a fixed level of symptoms. If variations in symptoms during infection – and their impacts on detectability – can be well characterised by epidemiologists and then included in predictive tools by modellers, decision makers will be able to make more informed choices as to which particular intervention, or combination of interventions, to pursue.

## DATA AVAILABILITY

The data used in our analyses are available in the supplementary files DataS1.csv and DataS2.csv. Analyses were performed in Matlab. Code is available for running the models, and is accessible at https://github.com/will-s-hart/EbolaVariableSymptoms.

## COMPETING INTERESTS

We have no competing interests.

## AUTHORS’ CONTRIBUTIONS

RNT conceived the research; All authors designed the study; RNT, LFRH and WSH carried out the research; RNT and WSH drafted the manuscript; All authors revised the manuscript and gave final approval for publication.

## FUNDING

RNT was funded by a Junior Research Fellowship from Christ Church, Oxford, and a JSPS Postdoctoral Research Fellowship (short-term, grant PE18029). WSH was funded by an EPSRC Excellence Award for his doctoral studies. LH was funded by an EPSRC undergraduate vacation bursary.

