## Supplementary Material for "Accurate forecasts of the effectiveness of interventions against Ebola may require models that account for variations in symptoms during infection"

### Supplementary Information

**AUTHORS**

W.S. Hart[1], L.F.R. Hochfilzer[1], N.J. Cunniffe[2], H. Lee[3], H. Nishiura[3], R.N. Thompson[1,4,5,*]

**AFFILIATIONS**

[1]Mathematical Institute, University of Oxford, Andrew Wiles Building, Radcliffe Observatory Quarter, Woodstock Road, Oxford OX2 6GG, UK

[2]Department of Plant Sciences, University of Cambridge, Downing Street, Cambridge CB2 3EA, UK

[3]Graduate School of Medicine, Hokkaido University, Hokkaido, Japan

[4]Department of Zoology, University of Oxford, South Parks Road, Oxford OX1 3PS, UK

[5]Christ Church, University of Oxford, St Aldates, Oxford OX1 1DP, UK

### Dependence of the detection rate on surveillance

In the main text, to illustrate the effects of different surveillance levels on the dynamics of the models, we considered a simple situation in which each host is surveyed for infection every $\Delta$ days. We assume that, when monitoring occurs, individuals in each of the three infectious compartments ($I_1, I_2$ and $I_3$, corresponding to the non-specific symptoms, gastrointestinal and deterioration phases of infection, respectively) have probabilities of successful detection and isolation given by $p_1, p_2$ and $p_3$. The dependence of the rates at which hosts are detected/isolated ($\delta_1, \delta_2$ and $\delta_3$) on the parameters $\Delta$, $p_1, p_2$ and $p_3$ can then be derived as follows.

If a host enters class $I_i$ (for *i* = 1,2 or 3) at a time that is uniformly distributed between surveillance times, then the expected time until the next surveillance time is $\Delta/2$ days. Ignoring progression through the infectious classes or to the *R* class, the expected number of surveys until the host is detected is given by $1/p_i$, so that the expected time after the first surveillance time until detection is $\Delta(1/p_i - 1)$ days. Summing these times gives

$$\text{Expected time in } I_i \text{ until detection} = \frac{\Delta}{2} + \Delta\left(\frac{1}{p_1} - 1\right),$$

$$= \Delta\left(\frac{1}{p_i} - \frac{1}{2}\right).$$

The rate of detection of a host in compartment $I_i$ is then given by the inverse of this expression, namely the equation in the main text.

### Robustness of results to parameter values used

Sampling frequency

The sampling frequency is characterised by the parameter Δ, which represents the time interval between surveillance rounds. We consider the robustness of the results in the main text to the sampling interval assumed when the models are fitted to the epidemic data (the model fitting stage), as well as to the sampling interval assumed when surveillance is altered in the fitted models to predict the effects of changing interventions (the intervention testing stage). For ease of notation, here we denote the sampling interval assumed during model fitting as $\Delta_1$ and the sampling interval under intervention testing as $\Delta_2$.

First, we assumed that the sampling interval during model fitting was $\Delta_1$ = 7 days, rather than the value of $\Delta_1$ = 21 days used in the main text. In Fig S1A, we show the model fits to data from the ongoing Ebola epidemic in the Democratic Republic of Congo, and in

Fig S1B, we display the model fits to the data from Liberia from the 2014-16 epidemic. In both cases, we find that the constant symptoms and variable symptoms models can replicate the data closely. When the surveillance interval is then changed, like in the main text we find that the predictions of the different models diverge (Fig S2).

We find similar results for $\Delta_1$ = 14 days (Figs S3 and S4), $\Delta_1$ = 21 days (Figs S5 – the model fitting is not shown in this case since it is given in Figs 3B and 3E of the main text) and $\Delta_1$ = 28 days (Figs S6 and S7). When predictions are made of the effect of intensifying surveillance, the constant symptoms model underestimates the total number of cases, whereas when predictions are made about the effects of relaxing surveillance then the constant symptoms model overestimates the total number of cases (for example, *cf.* Figs S4A and S4B).

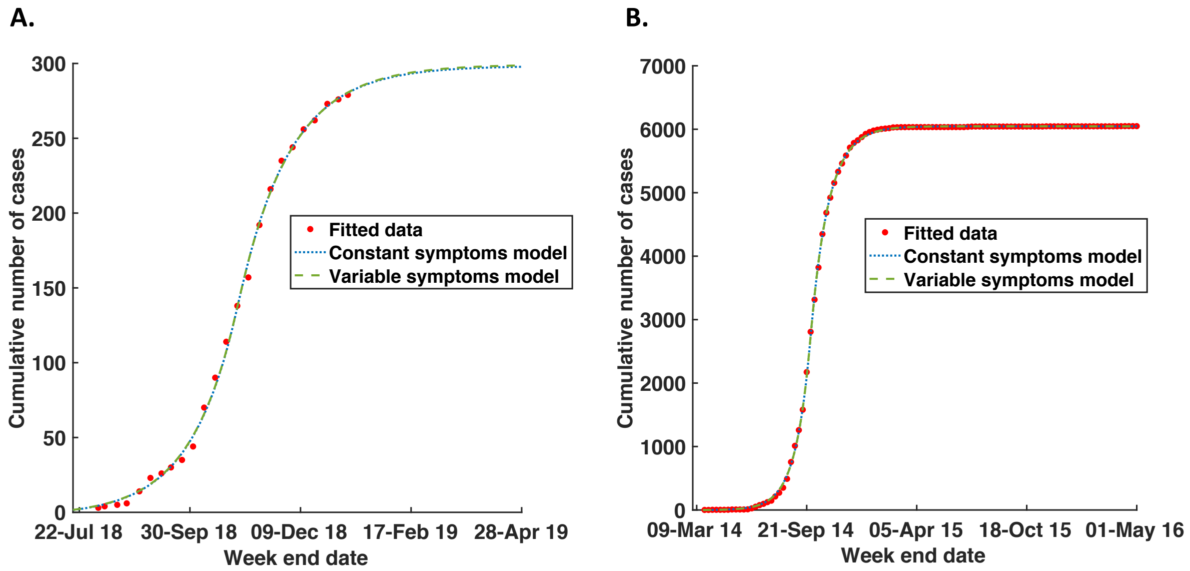


Figure S1. Model fits when the surveillance frequency is assumed to be $\Delta_1$ = 7 days. A. Model fits to the data from the ongoing epidemic in Beni and Kalunguta health zones in the Democratic Republic of Congo (red stars) for the constant symptoms (blue dotted) and variable symptoms (green dash) models. B. Model fits to the data from the 2014-16 Ebola epidemic in Liberia (red stars) for the constant symptoms (blue dotted) and variable symptoms (green dash) models. In panel A, for the constant symptoms model, fitted parameter values are: $\beta_0 = 1.19 \times 10^{-6}$ day$^{-1}$, $\beta_1 = 5.69 \times 10^{-7}$ day$^{-1}$, $T = 27/10/18$, $T_0 = 30/06/18$. In panel A, for the variable symptoms model, fitted parameter values are: $\beta_0 = 1.02 \times 10^{-6}$ day$^{-1}$, $\beta_1 = 4.82 \times 10^{-7}$ day$^{-1}$, $T = 26/10/18$, $T_0 = 28/06/18$. Other model parameters used in panel A are identical to those in Table 1 of the main text. In panel B, for the constant symptoms model, fitted parameter values are: $\beta_0 = 6.30 \times 10^{-8}$ day$^{-1}$, $\beta_1 = 2.91 \times 10^{-8}$ day$^{-1}$, $T = 24/09/14$, $T_0 = 21/03/14$. In panel B, for the variable symptoms model, fitted parameter values are: $\beta_0 = 5.41 \times 10^{-8}$ day$^{-1}$, $\beta_1 = 2.44 \times 10^{-8}$ day$^{-1}$, $T = 23/09/14$, $T_0 = 20/03/14$ for the variable symptoms model. Other model parameters used in panel B are identical to those in Table 2 of the main text.

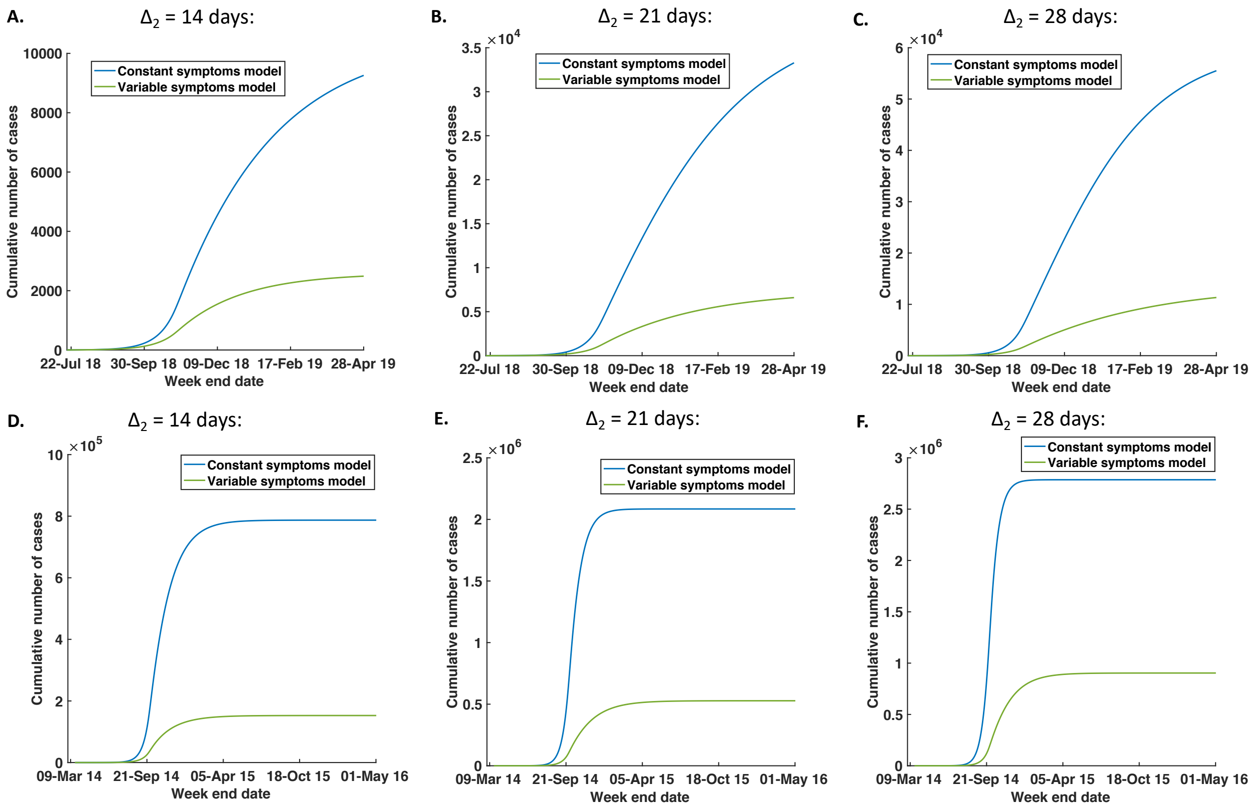


Figure S2. Using the constant symptoms and variable symptoms models to predict the effects of alternative interventions, after model fitting with $\Delta_1$ = 7 days (Fig S1). A. $\Delta_2$ = 14 days, using the constant symptoms model (blue) and variable symptoms model (green), for the model fitted to data from the ongoing epidemic in the Democratic Republic of Congo. B. Equivalent figure to A, with $\Delta_2$ = 21 days. C. Equivalent figure to A, with $\Delta_2$ = 28 days. D-F. Equivalent figures to A-C, using the data from the 2014-16 Ebola epidemic in Liberia. Values of other parameters are stated in the caption to Fig S1.

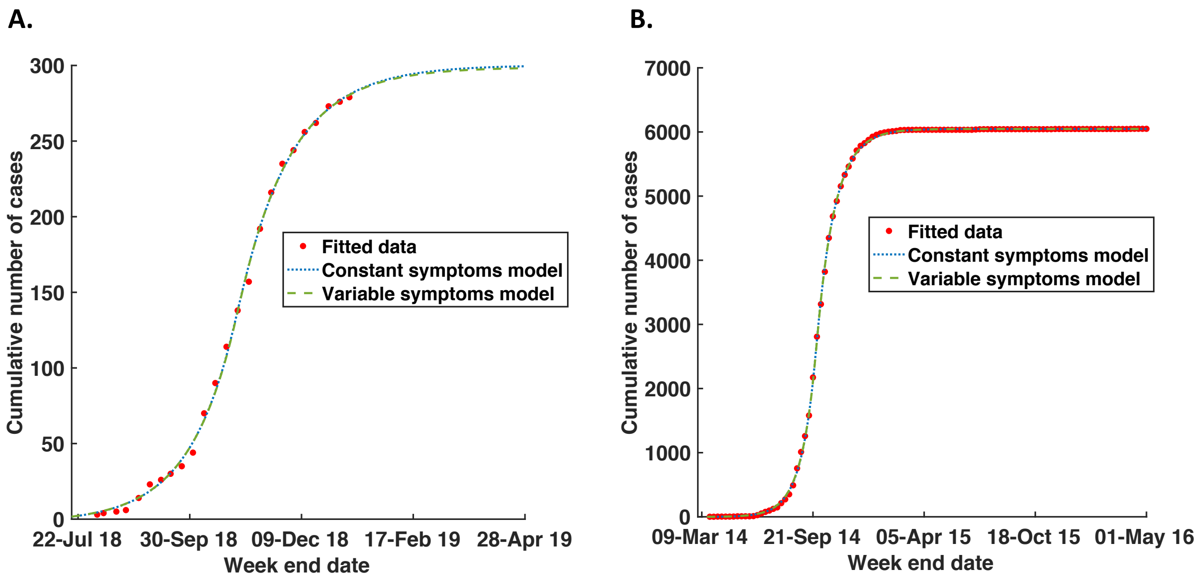


Figure S3. Model fits when the surveillance frequency is assumed to be $\Delta_1$ = 14 days. A. Model fits to the data from the ongoing epidemic in Beni and Kalunguta health zones in the Democratic Republic of Congo (red stars) for the constant symptoms (blue dotted) and variable symptoms (green dash) models. B. Model fits to the data from the 2014-16 Ebola epidemic in Liberia (red stars) for the constant symptoms (blue dotted) and variable symptoms (green dash) models. In panel A, for the constant symptoms model, fitted parameter values are: $\beta_0 = 9.29 \times 10^{-7}$ day$^{-1}$, $\beta_1 = 4.22 \times 10^{-7}$ day$^{-1}$, $T = 25/10/18$, $T_0 = 27/06/18$. In panel A, for the variable symptoms model, fitted parameter values are: $\beta_0 = 8.66 \times 10^{-7}$ day$^{-1}$, $\beta_1 = 3.88 \times 10^{-7}$ day$^{-1}$, $T = 25/10/18$, $T_0 = 26/06/18$. Other model parameters used in panel A are identical to those in Table 1 of the main text. In panel B, for the constant symptoms model, fitted parameter values are: $\beta_0 = 4.96 \times 10^{-8}$ day$^{-1}$, $\beta_1 = 2.13 \times 10^{-8}$ day$^{-1}$, $T = 22/09/14$, $T_0 = 21/03/14$. In panel B, for the variable symptoms model, fitted parameter values are: $\beta_0 = 4.61 \times 10^{-8}$ day$^{-1}$, $\beta_1 = 1.98 \times 10^{-8}$ day$^{-1}$, $T = 22/09/14$, $T_0 = 18/03/14$ for the variable symptoms model. Other model parameters used in panel B are identical to those in Table 2 of the main text.

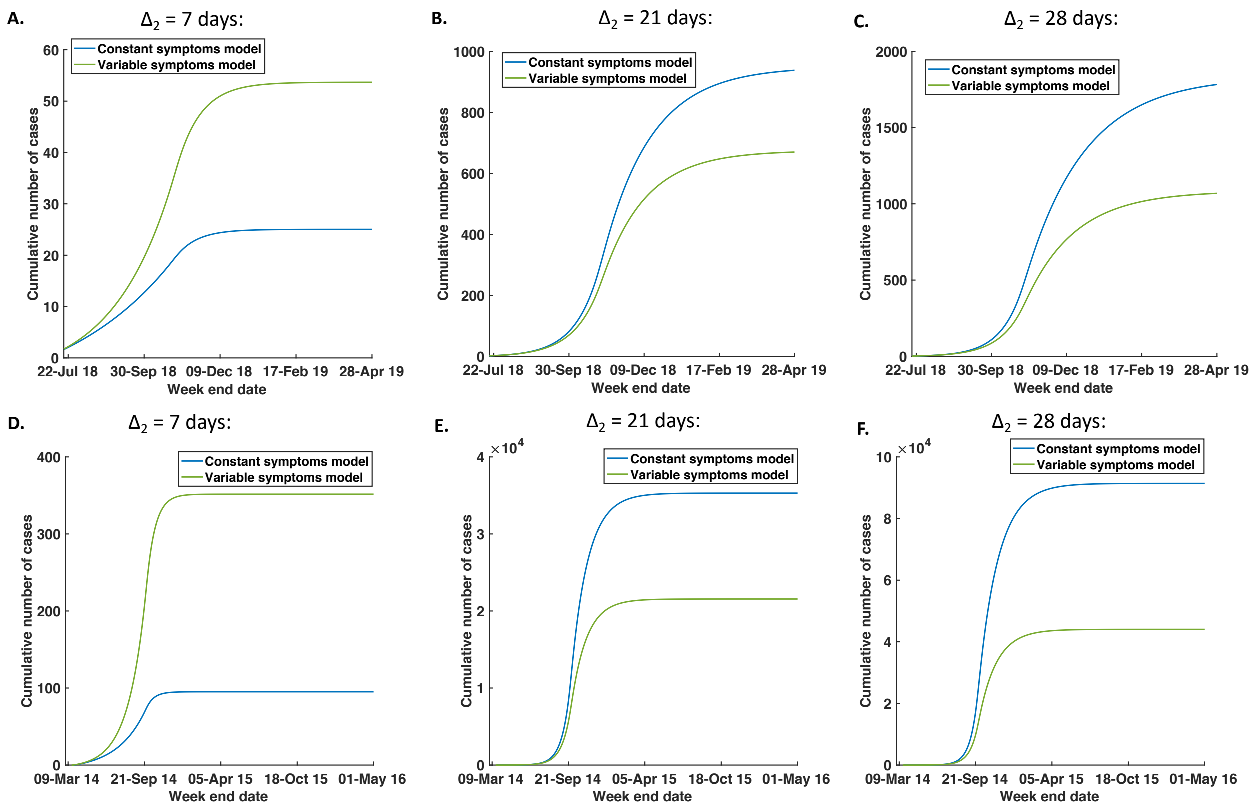


Figure S4. Using the constant symptoms and variable symptoms models to predict the effects of alternative interventions, after model fitting with $\Delta_1$ = 14 days (Fig S3). A. $\Delta_2$ = 7 days, using the constant symptoms model (blue) and variable symptoms model (green), for the model fitted to data from the ongoing epidemic in the Democratic Republic of Congo. B. Equivalent figure to A, with $\Delta_2$ = 21 days. C. Equivalent figure to A, with $\Delta_2$ = 28 days. D-F. Equivalent figures to A-C, using the data from the 2014-16 Ebola epidemic in Liberia. Values of other parameters are stated in the caption to Fig S3.

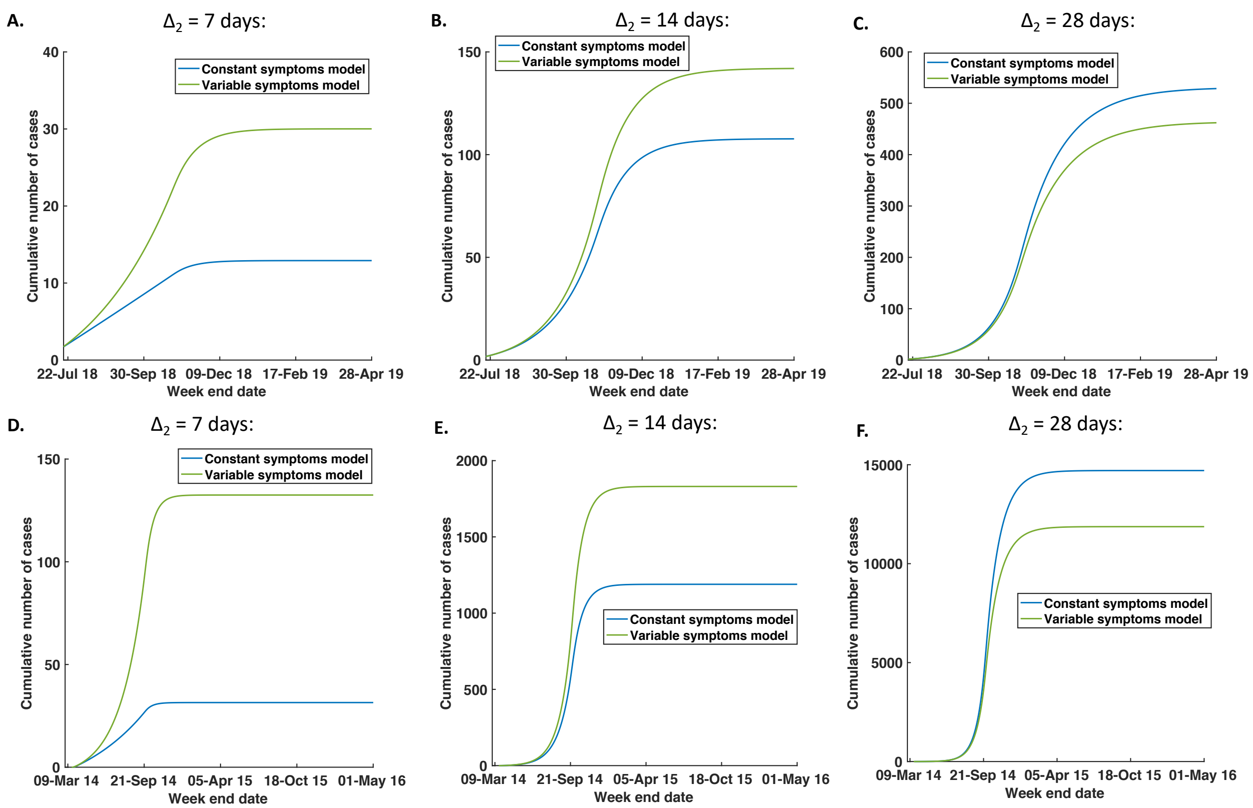


Figure S5. Using the constant symptoms and variable symptoms models to predict the effects of alternative interventions, after model fitting with $\Delta_1$ = 21 days (Figs 3B and 3E in the main text). A. $\Delta_2$ = 7 days, using the constant symptoms model (blue) and variable symptoms model (green), for the model fitted to data from the ongoing epidemic in the Democratic Republic of Congo. B. Equivalent figure to A, with $\Delta_2$ = 14 days. C. Equivalent figure to A, with $\Delta_2$ = 28 days. D-F. Equivalent figures to A-C, using the data from the 2014-16 Ebola epidemic in Liberia. Values of other parameters are stated in Tables 1 and 2 of the main text.

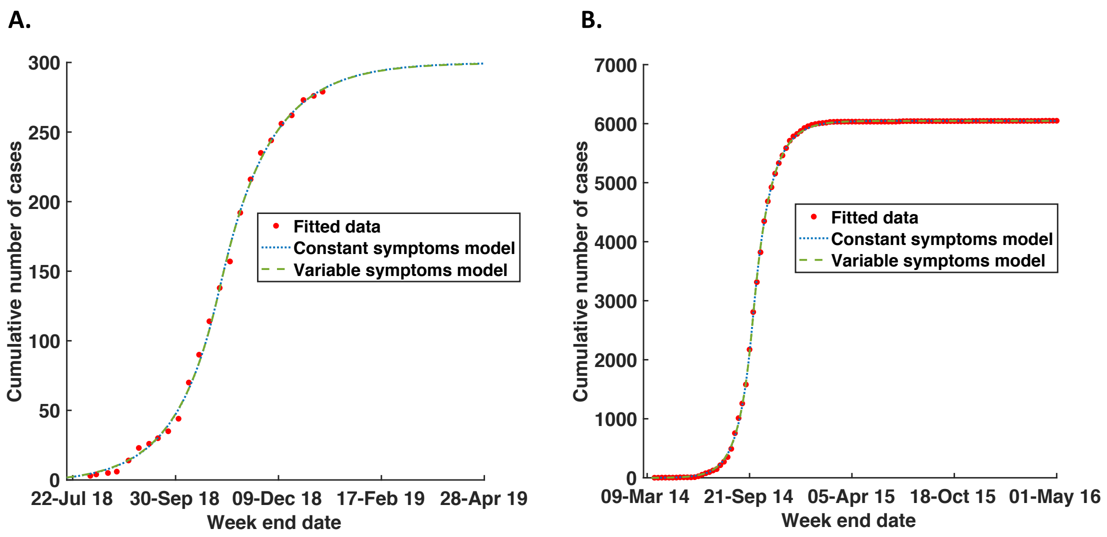


Figure S6. Model fits when the surveillance frequency is assumed to be $\Delta_1$ = 28 days. A. Model fits to the data from the ongoing epidemic in Beni and Kalunguta health zones in the Democratic Republic of Congo (red stars) for the constant symptoms (blue dotted) and variable symptoms (green dash) models. B. Model fits to the data from the 2014-16 Ebola epidemic in Liberia (red stars) for the constant symptoms (blue dotted) and variable symptoms (green dash) models. In panel A, for the constant symptoms model, fitted parameter values are: $\beta_0 = 8.05 \times 10^{-7}$ day$^{-1}$, $\beta_1 = 3.49 \times 10^{-7}$ day$^{-1}$, $T = 24/10/18$, $T_0 = 25/06/18$. In panel A, for the variable symptoms model, fitted parameter values are: $\beta_0 = 7.79 \times 10^{-7}$ day$^{-1}$, $\beta_1 = 3.38 \times 10^{-7}$ day$^{-1}$, $T = 24/10/18$, $T_0 = 24/06/18$. Other model parameters used in panel A are identical to those in Table 1 of the main text. In panel B, for the constant symptoms model, fitted parameter values are: $\beta_0 = 4.30 \times 10^{-8}$ day$^{-1}$, $\beta_1 = 1.76 \times 10^{-8}$ day$^{-1}$, $T = 21/09/14$, $T_0 = 19/03/14$. In panel B, for the variable symptoms model, fitted parameter values are: $\beta_0 = 4.16 \times 10^{-8}$ day$^{-1}$, $\beta_1 = 1.71 \times 10^{-8}$ day$^{-1}$, $T = 21/09/14$, $T_0 = 17/03/14$ for the variable symptoms model. Other model parameters used in panel B are identical to those in Table 2 of the main text.

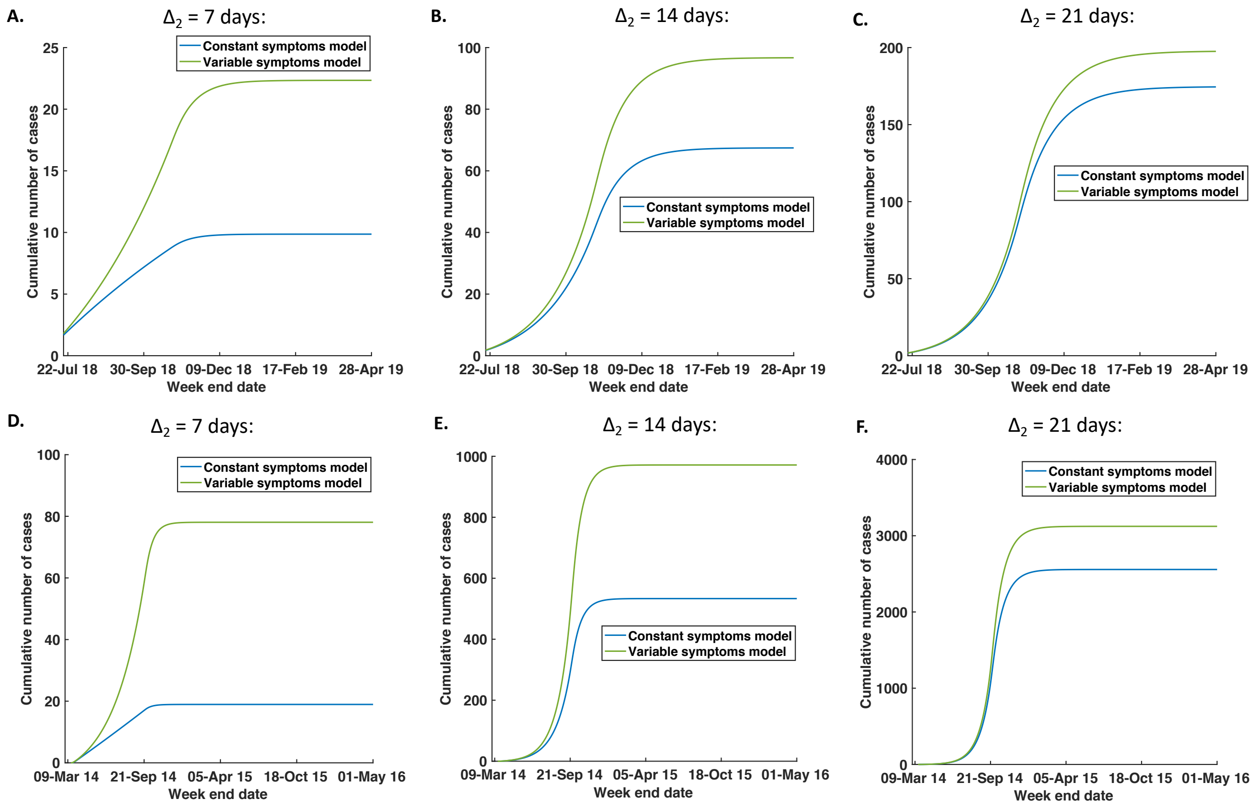


Figure S7. Using the constant symptoms and variable symptoms models to predict the effects of alternative interventions, after model fitting with $\Delta_1$ = 28 days (Fig S6). A. $\Delta_2$ = 7 days, using the constant symptoms model (blue) and variable symptoms model (green), for the model fitted to data from the ongoing epidemic in the Democratic Republic of Congo. B. Equivalent figure to A, with $\Delta_2$ = 14 days. C. Equivalent figure to A, with $\Delta_2$ = 21 days. D-F. Equivalent figures to A-C, using the data from the 2014-16 Ebola epidemic in Liberia. Values of other parameters are stated in the caption of Fig S6.

### Detection probability

In our analyses in the main text, in the constant symptoms model we assumed that the detection probability when a host was sampled was $p_1 = p_2 = p_3 = 0.6$. In the variable symptoms model, we assumed that the detection probabilities were $p_1 = 0.1$, $p_2 = 0.8$ and $p_3 = 0.9$ in early, mid and late infection, respectively. The mean value of $p_1, p_2$ and $p_3$ was therefore the same in both models.

Here we consider the robustness of our results to the precise values of $p_1$, $p_2$ and $p_3$ used. In particular, we perform two separate analyses. In the first, the value of the detection probability in the constant symptoms model is varied (with $p_2$ in the variable symptoms model also varied so that the mean detection probability in both models remains equal). In the second, the variation of $p_1$, $p_2$ and $p_3$ in the variable symptoms model about the mean value is changed. The results of the first analysis are shown in Fig S8, and the results of the second analysis are shown in Fig S9. We do not display the model fits, since these are identical to those in Figs 3B and 3E of the main text when assessed by eye, and instead show the predictions of the fitted models when interventions are changed. In both Figs S8 and S9, we find that results that are qualitatively identical to those in the main text, namely that the constant symptoms model leads to an underestimation of the number of cases under intensified surveillance compared to the more realistic variable symptoms model.

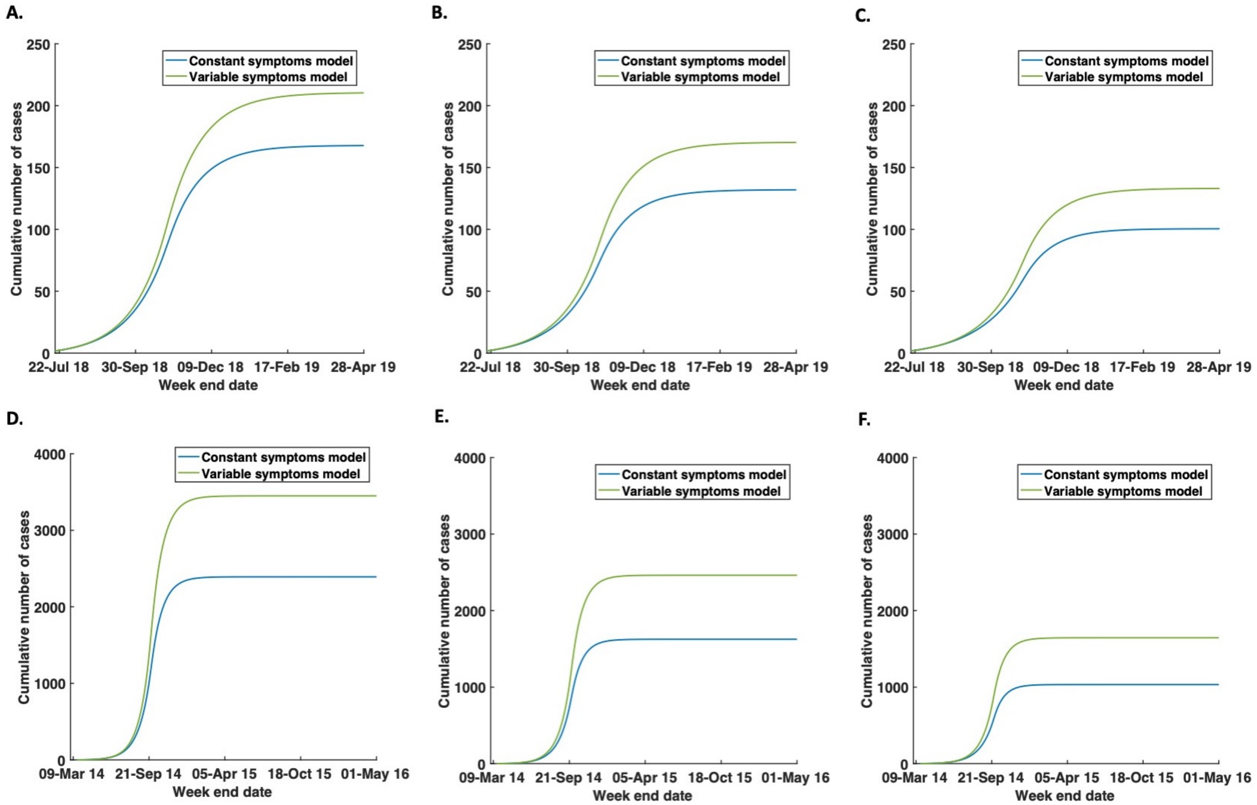


Figure S8. Using the constant symptoms and variable symptoms models to predict the effects of intensified surveillance, for different assumed mean detection probabilities. A. Low surveillance (mean detection probability of 0.367), using the constant symptoms model (blue) and variable symptoms model (green), for the model fitted to data from the ongoing epidemic in the Democratic Republic of Congo. B. Equivalent figure to A but for medium surveillance (mean detection probability of 0.5). C. Equivalent figure to A but for high surveillance (mean detection probability of 0.663). D-F. Equivalent figures to A-C, using data from the 2014-16 Ebola epidemic in Liberia. In the variable symptoms model, the values of $p_1$ and $p_3$ are held fixed at 0.1 and 0.9, respectively, with $p_2$ varied so that the mean value of $p_1$ $p_2$ and $p_3$ is equal to the mean detection probability. This corresponds to values of $p_2$ in the variable symptoms model of 0.1 (panels A and D), 0.5 (panels B and E) and 0.9 (panels C and F), respectively. Other parameters values are given in Tables 1 and 2 of the main text.

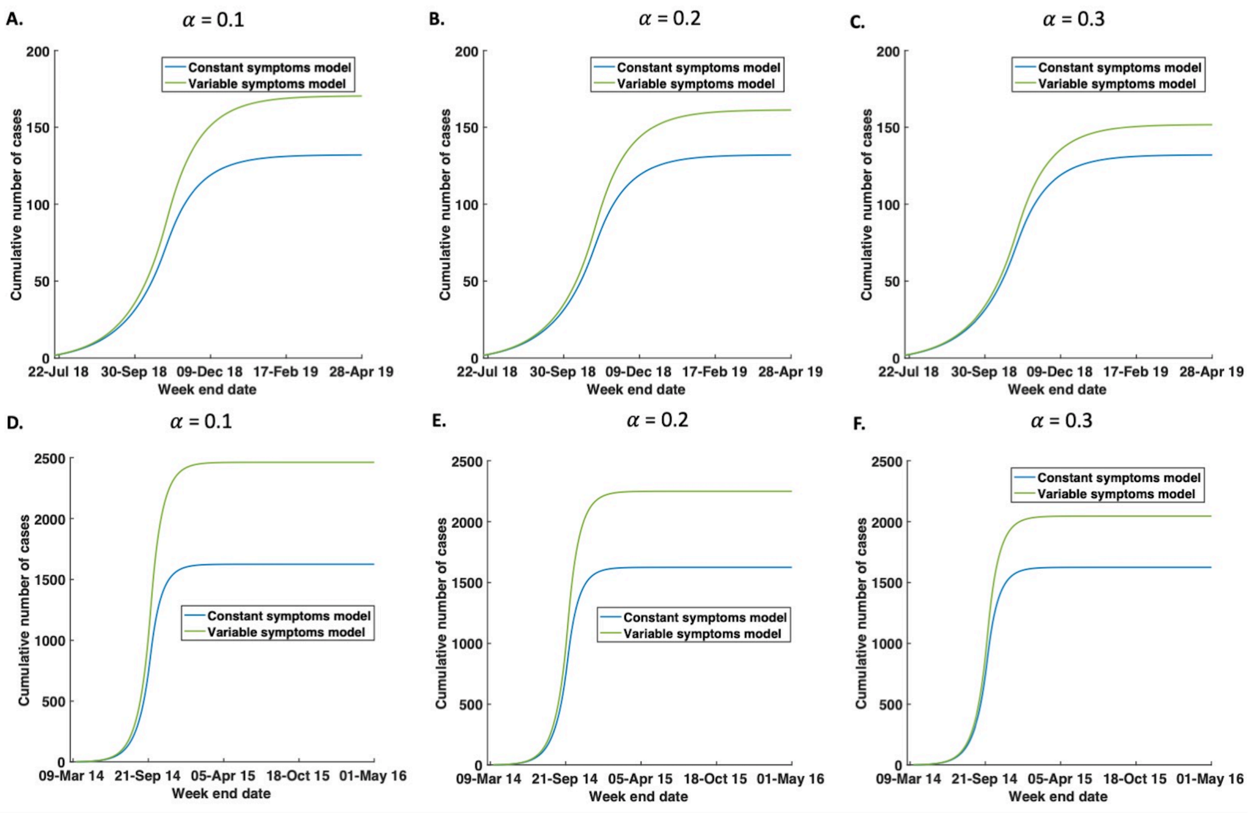


Figure S9. Using the constant symptoms and variable symptoms models to predict the effects of intensified surveillance, for different variations in symptom expression during infection. In this analysis, we fix the detection probability in the constant symptoms model to $p_1 = p_2 = p_3 = 0.5$, and take the value $p_2 = 0.5$ in the variable symptoms model. In the variable symptoms model, we then take $p_1 = \alpha$ and $p_3 = (1 - \alpha)$, for different values of the parameter $\alpha$. A. $\alpha = 0.1$, using the constant symptoms model (blue) and variable symptoms model (green), for the model fitted to data from the ongoing epidemic in the Democratic Republic of Congo. B. Equivalent figure to A, with $\alpha = 0.2$. C. Equivalent figure to A, with $\alpha = 0.3$. D-F. Equivalent figures to A-C, using the data from the 2014-16 Ebola epidemic in Liberia. Other parameters values are given in Tables 1 and 2 of the main text.

### The timing of introduction in surveillance

In Fig 3 of the main text, we used the constant symptoms and variable symptoms models to predict the impact of intensified surveillance on the dynamics of an Ebola epidemic. Implicit in that analysis was that the increased level of surveillance was in place throughout the epidemic. However, in outbreak response settings, an intensification of surveillance could occur after the epidemic is underway. Here, we consider a scenario in which surveillance is instead intensified at a date $\tau$, after the first observed cases in the epidemic data. We consider $\tau$ dates of 23$^{rd}$ August 2018, 12$^{th}$ September 2018 and 2$^{nd}$ October 2018 for the epidemic in the Democratic Republic of Congo, and 2$^{nd}$ May 2014, 11$^{th}$ June 2014 and 21$^{st}$ July 2014 for the epidemic in Liberia. All parameters are as in Table 1 and Table 2 of the main text.

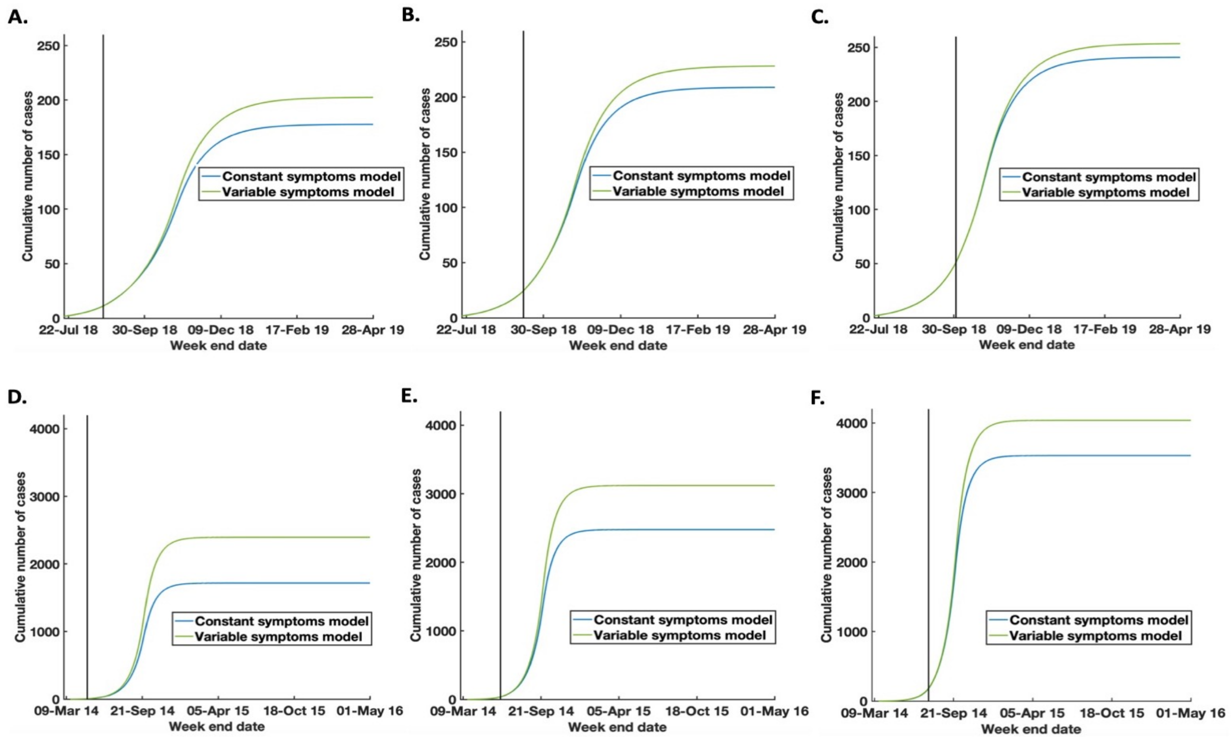


Figure S10. Using the constant symptoms and variable symptoms models to predict the effects of intensified surveillance, introduced at different dates ($\tau$) during an Ebola epidemic. A. $\tau =$ 23-Aug 18

using the constant symptoms model (blue) and variable symptoms model (green), for the model fitted to data from the ongoing epidemic in the Democratic Republic of Congo. B. $\tau$ = 12-Sep 18 using the constant symptoms model (blue) and variable symptoms model (green), for the model fitted to data from the ongoing epidemic in the Democratic Republic of Congo. C. $\tau$ = 2-Oct 18 using the constant symptoms model (blue) and variable symptoms model (green), for the model fitted to data from the ongoing epidemic in the Democratic Republic of Congo. D. $\tau$ = 2-May 14 using the constant symptoms model (blue) and variable symptoms model (green), for the model fitted to data from the 2014-16 epidemic in Liberia. E. $\tau$ = 11-Jun 14 using the constant symptoms model (blue) and variable symptoms model (green), for the model fitted to data from the 2014-16 epidemic in Liberia. F. $\tau$ = 21-Jul 14 using the constant symptoms model (blue) and variable symptoms model (green), for the model fitted to data from the 2014-16 epidemic in Liberia. Other parameters values are given in Tables 1 and 2 of the main text. The black vertical lines indicate the times at which surveillance is intensified.

### Model for time-dependent infectiousness

In Figs 2-4 of the main text, to isolate the effect of variable symptoms alone on epidemic predictions, we assumed that the infection rate was constant for all infectious hosts. However, in reality the infectiousness of a host is likely to vary throughout an Ebola infection too. We therefore considered an additional analysis (Fig 5 of the main text), in which infectiousness was assumed to vary during an infection. Here we described the model used to perform that analysis, and its parameterisation.

We extend the $SEI_1I_2I_3RC$ model so that individuals in compartment $I_i$ have infection rate $\beta^{(i)}(t)$ (for *i* = 1, 2 or 3). This leads to the extended model

$$\frac{\mathrm{d}S}{\mathrm{d}t} = -S\left(\beta^{(1)}(t)I_1 + \beta^{(2)}(t)I_2 + \beta^{(3)}(t)I_3\right),$$

$$\frac{\mathrm{d}E}{\mathrm{d}t} = S\left(\beta^{(1)}(t)I_1 + \beta^{(2)}(t)I_2 + \beta^{(3)}(t)I_3\right) - \gamma E,$$

$$\frac{\mathrm{d}I_1}{\mathrm{d}t} = \gamma E - \mu I_1 - \delta_1 I_1,$$

$$\frac{\mathrm{d}I_2}{\mathrm{d}t} = \mu I_1 - \mu I_2 - \delta_2 I_2,$$

$$\frac{dI_3}{dt} = \mu I_2 - \mu I_3 - \delta_3 I_3,$$

$$\frac{dR}{dt} = \mu I_3,$$

$$\frac{dC}{dt} = \delta_1 I_1 + \delta_2 I_2 + \delta_3 I_3.$$

We consider three particular forms of this model: i) the constant infectiousness model in which $\beta^{(1)}(t) = \beta^{(2)}(t) = \beta^{(3)}(t) = \beta(t)$, i.e. the model underlying Figs 2-4 of the main text; ii) an increasing infectiousness model in which we assume $\beta^{(1)}(t) = 0.5\beta(t)$, $\beta^{(2)}(t) = \beta(t)$ and $\beta^{(3)}(t) = 1.5\beta(t)$; iii) a decreasing infectiousness model in which $\beta^{(1)}(t) = 1.5\beta(t)$, $\beta^{(2)}(t) = \beta(t)$ and $\beta^{(3)}(t) = 0.5\beta(t)$. In each case, we still assume that the infection rate of all infectious individuals is reduced at some time $T$ during the epidemic, so that

$$\beta(t) = \begin{cases} \beta_0 \text{ for } t \leq T \text{ days,} \\ \beta_1 \text{ for } t > T \text{ days.} \end{cases}$$

The values of the parameters $\beta_0$, $\beta_1$ and $T$, in addition to the epidemic start date $T_0$, were fitted to the epidemic data under consideration, either under the assumption of constant symptoms ($\delta_1 = \delta_2 = \delta_3 = \delta$) or variable symptoms ($\delta_1 < \delta_2 < \delta_3$). All other parameter values are given in Table 1 and Table 2 of the main text.

**Introducing additional epidemiological complexity in the models**

To illustrate the principle that models with variable symptoms can generate different predictions compared to models with constant symptoms, we considered the simplest possible extension of the SEIR model in which the level of symptoms increases throughout infection. Here, we illustrate how additional realism could be included in such a model.

In particular, we consider an extension of the $SEI_1I_2I_3RC$ model, in which we assume that some individuals recover before they progress to the deterioration phase. In reality, an individual progressing to the deterioration phase is very likely to die from the Ebola infection. In the absence of surveillance, we assume a proportion, $q$, of hosts recover during the gastrointestinal phase of infection (represented by the $I_2$ compartment), while the remainder continue to the deterioration phase (the $I_3$ compartment) and subsequently die. This leads to the extended model

$$\frac{\mathrm{d}S}{\mathrm{d}t} = -\beta(t)S(I_1 + I_2 + I_3),$$

$$\frac{\mathrm{d}E}{\mathrm{d}t} = \beta(t)S(I_1 + I_2 + I_3), - \gamma E,$$

$$\frac{\mathrm{d}I_1}{\mathrm{d}t} = \gamma E - \mu I_1 - \delta_1 I_1,$$

$$\frac{\mathrm{d}I_2}{\mathrm{d}t} = \mu I_1 - \mu I_2 - \delta_2 I_2,$$

$$\frac{\mathrm{d}I_3}{\mathrm{d}t} = (1 - q)\mu I_2 - \mu I_3 - \delta_3 I_3,$$

$$\frac{\mathrm{d}R}{\mathrm{d}t} = q\mu I_2,$$

$$\frac{\mathrm{d}D}{\mathrm{d}t} = \mu I_3,$$

$$\frac{\mathrm{d}C}{\mathrm{d}t} = \delta_1 I_1 + \delta_2 I_2 + \delta_3 I_3.$$

In this new $SEI_1I_2I_3RDC$ model, we differentiate explicitly between hosts that have recovered without being detected ($R$) and those that are detected at death ($D$).

As before, we estimated the parameters $\beta_0$, $\beta_1$, $T$ and $T_0$ (defined analogously to the corresponding parameters in the baseline model) by fitting the model to the Beni and Liberia datasets, under the assumptions of constant symptoms and variable symptoms in turn. When fitting the model, we assumed that the number of observed cases corresponds to the sum of $C$ and $D$ in the model. The values of other parameters are as in Table 1 and Table 2 of the main text. In each case, the model fits are identical to those in Figs 3B and 3E of the main text when assessed by eye, and are therefore not shown.

We find that, when a high proportion of individuals recover rather than being detected, predictions of the constant symptoms and variable symptoms models differ by more than when a low proportion of individuals recover rather than being detected. However, in every case that we considered, there is a substantial difference between the predicted numbers of cases of the models, supporting our assertion that it may be necessary to include variable symptoms during infection in models of Ebola intervention strategies.

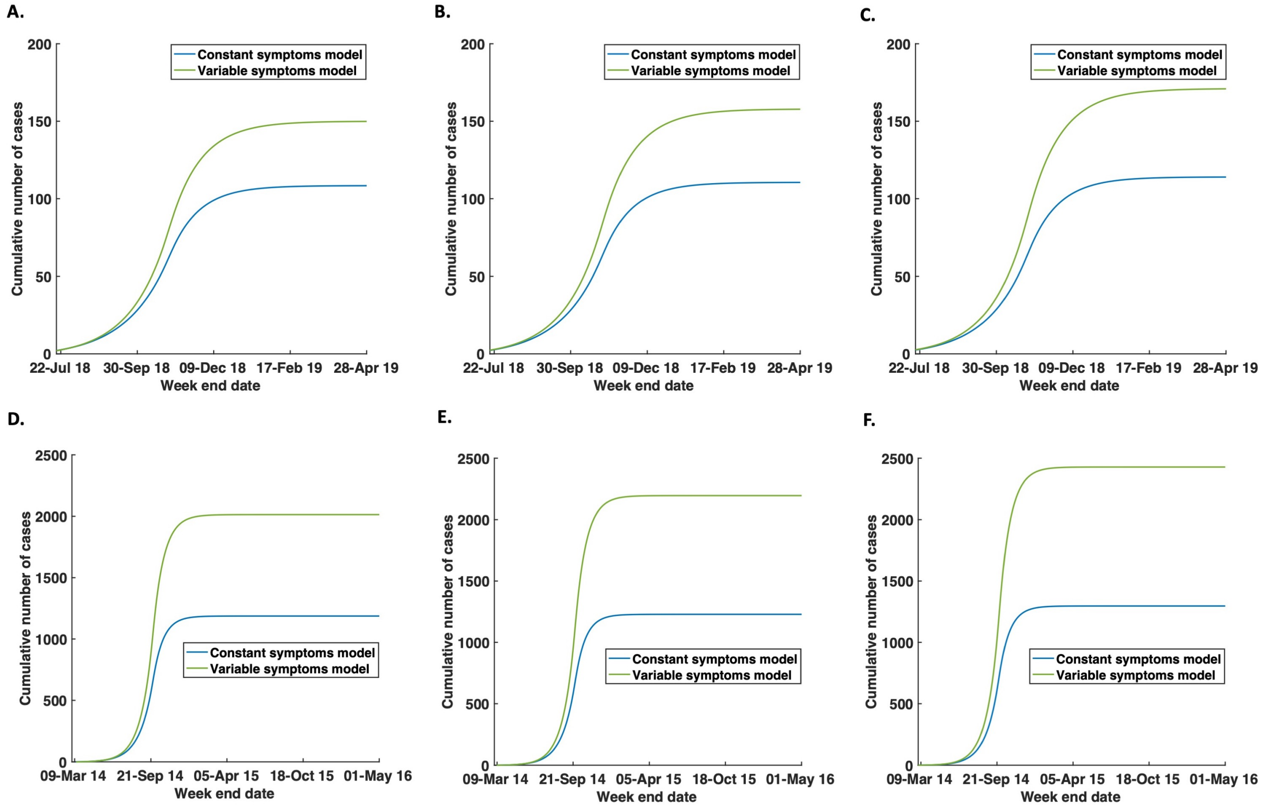


Figure S11. Using the constant symptoms and variable symptoms models to predict the effects of intensified surveillance, when some hosts recover prior to the deterioration phase of an Ebola infection. The proportion of hosts recovering rather than passing into the $I_3$ class is denoted by $q$. A. $q = 0.3$, using the constant symptoms model (blue) and variable symptoms model (green), for the model fitted to data from the ongoing epidemic in the Democratic Republic of Congo. B. Equivalent to A, but with $q = 0.5$. C.

Equivalent to A, but with $q = 0.7$. D-F. Equivalent figures to A-C, using the data from the 2014-16 Ebola epidemic in Liberia. Other parameters values are given in Tables 1 and 2 of the main text.
